# Inhibitory fear memory engram in the mouse central lateral amygdala

**DOI:** 10.1101/2023.11.30.565632

**Authors:** Wen-Hsien Hou, Meet Jariwala, Kai-Yi Wang, Anna Seewald, Yu-Ling Lin, Alessia Ricci, Francesco Ferraguti, Cheng-Chang Lien, Marco Capogna

**Author notes:** Deceased (Marco Capogna). Both authors equally contributed (Wen-Hsien Hou, Meet Jariwala).

## Abstract

Engrams are cellular substrates of memory traces that have been identified in various brain areas, including the amygdala. Most engrams identified so far are formed by excitatory, glutamatergic neurons. However, little attention has been paid to defining GABAergic inhibitory engrams. Here, we report an inhibitory engram in the central lateral amygdala (CeL), a crucial area for Pavlovian fear conditioning. This engram is primarily composed of GABAergic somatostatin-expressing (SST+) and to a lesser extent of protein kinase C-δ-expressing [PKC-δ(+)] neurons. Fear memory is accompanied by a preferential enhancement of mIPSC frequency onto PKC-δ(+) neurons as well as a general increment of amplitude. Moreover, non-engram cells exhibit higher mIPSC frequency than engram cells. The inhibition of the CeL GABAergic engram disinhibits the activity of engram-targeted areas and increases selectively the encoded fear expression. Our data defines the behavioral function of an engram formed exclusively by GABAergic inhibitory neurons in the mammalian CNS.

## INTRODUCTION

Engrams are groups of neurons that are believed to represent the neural substrates for memory storage and recall (Rao-Ruiz et al., 2019). The critical role of engrams on memories has been established through loss-of-function, gain-of-function, and mimicry experiments (Josselyn and Tonegawa, 2020). Engrams are found in many neuronal circuits, including the amygdala. For example, the elimination of engram neurons in the lateral amygdala (LA) impairs auditory fear memory retrieval (Han et al., 2009), optogenetic reactivation of a basolateral amygdala (BLA) engram enhances fear memory (Redondo et al., 2014), and the activation of a BLA engram promotes odor fear memories (Vetere et al., 2019).

Various types of GABAergic neurons play a key role in gating the output of principal neurons projecting outside the amygdala (Capogna, 2014; Krabbe et al., 2018). Therefore, it is likely that synaptic inhibition plays a determinant role in the formation of excitatory engrams. Several lines of evidence support this view, for instance inhibition of parvalbumin (PV)+ interneurons during fear conditioning (FC) increases the size of LA engram (Morrison et al., 2016). Silencing somatostatin (SST)+ interneurons during contextual FC enhances the size of hippocampal dentate granule cells engram (Stefanelli et al., 2016). A recent computational approach has suggested that inhibitory engrams are essential components for maintaining an excitatory-inhibitory balance during the formation and recall of associative memories (Barron et al., 2017). In addition, inhibitory engrams were suggested to have a distinct functional role compared to excitatory engrams (Koolschijn et al., 2019; Sun et al., 2020a).

One significant progress would be to characterize inhibitory engrams in brain circuits with clear-cut behavioral significance. The lateral division of the central amygdala (CeL) is an ideal microcircuit for this purpose, as it contains primarily GABAergic cells expressing either PKC-δ, SST (Ciocchi et al., 2010; Haubensak et al., 2010; Li et al., 2013), or corticotropin-releasing factor (CRF) (Fadok et al., 2017). PKC-δ(+) and SST+ neurons have been identified as CeL “fear-off” or CeL “fear-on” neurons, respectively, because they reduce or increase their firing after a conditioned stimulus (Ciocchi et al., 2010; Haubensak et al., 2010; Li et al., 2013; Yu et al., 2016). On the other hand, CRF+ neurons mediate conditioned flight (Fadok et al., 2017). Interestingly, these GABAergic neurons are extensively interconnected (Fadok et al., 2017; Hou et al., 2016; Hunt et al., 2017). Furthermore, SST+ neurons receive direct synaptic inputs from the LA and from the paraventricular nucleus of the thalamus, and both inputs are critical for auditory FC or the establishment of fear memory and the expression of fear responses (Li et al., 2013; Penzo et al., 2015). Conversely, PKC-δ(+) neurons inhibit CeM neurons to elicit defensive responses through connections to downstream targets (Ciocchi et al., 2010; Haubensak et al., 2010) and also project back to the LA to convey information about the unconditioned stimulus (US) during FC (Yu et al., 2017).

Therefore, given the importance of the CeL for fear memory processing, we hypothesized that inhibitory engrams are formed in this area during FC. In this study, we observed inhibitory plasticity in the CeL after FC and identified CeL engram cells projecting locally and to extra-amygdaloid areas. Inhibition of the CeL engram neurons facilitates fear memory expression. Our findings shed new light on the complex mechanisms involved in the formation and regulation of fear memories.

## RESULTS

### Fear memory retrieval is associated with inhibitory synaptic plasticity in the CeL

First, we tested whether after cued FC, fear memory retrieval was accompanied by changes in the strength of GABAergic transmission impinging on CeL neurons of the mouse amygdala. We split the experimental mice into two groups: the FC group, which underwent five pairings of a neutral auditory conditioned stimulus (CS) co-terminating with a foot shock (US), and the control group (Ctrl), which was exposed only to the CS (Figure 1A, left). The FC group exhibited a progressive increase in freezing, indicating successful acquisition of a conditioned response (Figure 1B). The mice from both groups were then subjected to a fear memory retrieval test 24 hours (hrs) later, consisting of the exposure to the tone CS (in a different context) in the absence of the US (Figure 1A, middle). The FC group exhibited significantly higher freezing levels than the Ctrl group in response to the CS (Figure 1B). Within 1.5 hrs from the fear memory retrieval test, we measured *ex vivo* spontaneous inhibitory postsynaptic currents (sIPSCs), in the presence of 2 mM kynurenic acid (KA) to block glutamatergic synaptic currents, by performing whole cell recordings of CeL neurons in acute brain slices prepared from both groups of mice (Figure 1A, right; Figure 1C). We found that the sIPSC frequency was significantly higher in CeL neurons recorded from the FC group than those from the Ctrl group, whereas the amplitude of the sIPSCs did not differ between the two groups (Figures 1D-F).

**Figure 1.**
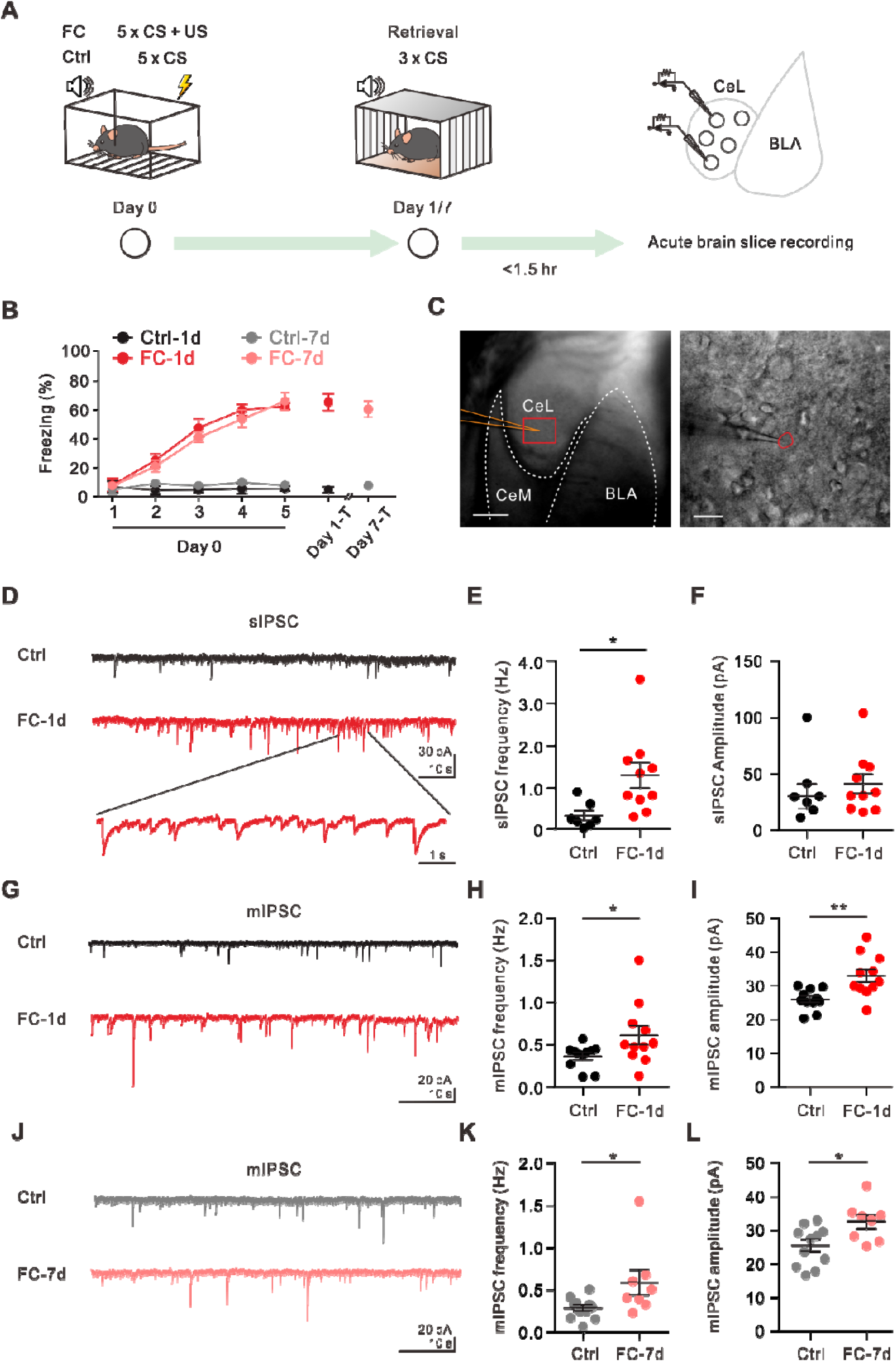
Fear conditioning leads to increased spontaneous inhibitory transmission onto CeL neurons. **(A)** Experimental paradigm. **(B)** Average freezing time of Ctrl (CS-only: Ctrl-1d, Ctrl-7d) and FC (CS+US: FC-1d, FC-7d) groups during auditory associative fear conditioning and after fear memory retrieval test (3xCS) on day 1 (Day 1-T) and day 7 (Day 7-T). p < 0.0001, two-way ANOVA. **(C)** Micrograph of the amygdaloid complex taken from a coronal section of an acute slice and diagram of the mouse CeL region in 10X (left) and 40X magnification (right). Scale bars, 200 μm (left) and 20 μm (right). **(D)** Example traces illustrating 60-s sweeps of sIPSCs recorded from control and fear conditioned mice (black and red, respectively), with the enlarged 6-s red trace shown below indicating the inward IPSCs recorded with a high [Cl^-^] internal solution. **(E)** sIPSC frequency (Ctrl, 0.32 ± 0.12 Hz, n = 7 cells; FC-1d, 1.27 ± 0.30 Hz, n = 10 cells; p = 0.042, Mann-Whitney test). *p < 0.05. **(F)** sIPSC amplitude (Ctrl, 46.58 ± 11.24 pA, n = 7 cells; FC-1d, 41.43 ± 8.46 pA, n = 10 cells; p = 0.35, Mann-Whitney test). **(G)** Example traces illustrating 60-s sweeps of mIPSCs recorded after 3xCS on day 1 from control and fear conditioned mice (black and red, respectively). **(H)** mIPSC frequency (Ctrl, 0.36 ± 0.04 Hz, n = 11 cells; FC-1d, 0.61 ± 0.11 Hz, n = 11 cells; p = 0.0321, Mann-Whitney test). *p < 0.05. **(I)** mIPSC amplitude (Ctrl, 25.98 ± 0.95 pA, n = 11 cells; FC-1d, 32.96 ± 1.85 pA, n = 11 cells; p = 0.0026, Mann-Whitney test). **p < 0.01. **(J)** Example traces illustrating a 60-s sweep of mIPSCs recorded after 3xCS on day 7 from Ctrl and FC mice (black and red, respectively). **(K)** mIPSC frequency (Ctrl, 0.29 ± 0.04 Hz, n = 11 cells; FC-7d, 0.59 ± 0.15 Hz, n = 8 cells; p = 0.015, Mann-Whitney test). *p < 0.05. **(L)** mIPSC amplitude (Ctrl, 25.48 ± 1.80 pA, n = 11 cells; FC-7d, 33.57 ± 2.02 pA, n = 8 cells; p = 0.0328, Mann-Whitney test). *p < 0.05.

To isolate activity-independent GABAergic neurotransmission, we also measured miniature inhibitory postsynaptic currents (mIPSCs) in the presence of 1 μM tetrodotoxin to block action potentials and 2 mM KA to block glutamatergic synaptic currents. We found that the mIPSC frequency and amplitude recorded from CeL neurons were significantly higher in the FC group than in the Ctrl group (Figures 1G-I). Fear memory can be long lasting (Alberini, 2005). To determine if the observed strengthening of GABAergic transmission persisted, we conducted fear memory retrieval tests 7 days after FC (Figure 1A, left). Indeed, we observed that the mIPSC frequency and amplitude remained significantly higher than that observed in the Ctrl group (Figures 1J-L). Taken together, these results demonstrate that FC strengthens inhibitory synaptic transmission onto CeL neurons at least up to 7 days.

### Fear memory is accompanied by inhibitory synaptic plasticity impinging mainly on CeL PKC-**δ(+)** GABAergic neurons

The CeL is composed of SST+ and PKC-δ(+) GABAergic neurons that are equally numerous and interconnected (Ciocchi et al., 2010; Haubensak et al., 2010; Shrestha et al., 2020). To determine whether inhibitory synaptic plasticity that accompanied fear memory retrieval is neuron-type specific, we used two complementary mouse transgenic lines to identify CeL neurons *ex vivo* (Figure 2A).

**Figure 2.**
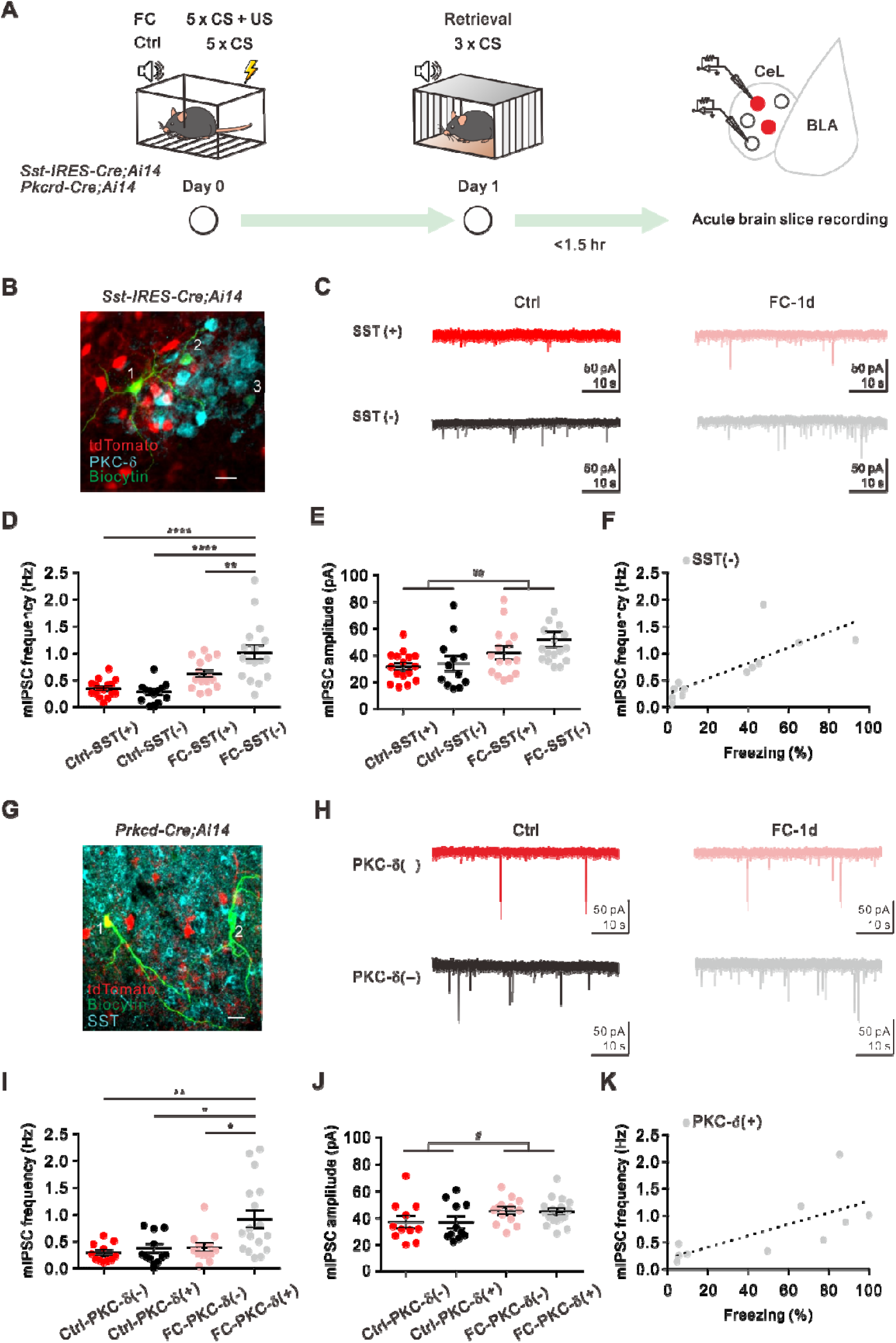
Fear conditioning strengthens mIPSCs mainly in SST(-)/PKC-δ(+) CeL neurons. **(A)** Experimental schematic and timeline. **(B)** The CeL section of a *Sst-IRES-Cre;Ai14* mouse with *post hoc* immunostaining of PKC-δ and biocytin. Confocal image indicated a CeL biocytin-filled, recorded a SST+/PKC-δ(-) neuron (#1) and two SST-/PKC-δ(+) neurons (#2 and #3). Scale bar, 20 μm. **(C)** Representative mIPSC recordings from SST+ (red) and SST− (black) neurons in the CeL of Ctrl and FC-1d groups. **(D)** mIPSC frequency of CeL neurons of Ctrl and FC-1d groups (SST+, Ctrl: 0.34 ± 0.05 Hz, n = 16 cells; FC-1d: 0.61 ± 0.07 Hz, n = 15 cells; SST−, Ctrl: 0.28 ± 0.07 Hz, n = 12 cells; FC-1d: 1.10 ± 0.13 Hz; n = 17 cells; *p < 0.05, ***p < 0.001, ****p < 0.0005, two-way ANOVA with Tukey’s multiple comparisons test). **(E)** mIPSC amplitude of CeL neurons of Ctrl and FC-1d groups (SST+, Ctrl: 31.81 ± 2.71 pA, n = 16 cells; FC-1d: 42.07 ± 4.64 pA, n = 15 cells; SST−, Ctrl: 34.14 ± 5.53 pA, n = 12 cells; FC-1d: 52.03 ± 5.56 pA; n = 17 cells. main treatment effect: Ctrl vs. FC, ^##^p = 0.0046, two-way ANOVA). **(F)** Plot of mIPSC frequency of SST-neurons against animal freezing levels (n = 12 animals, Linear regression R square = 0.627, p = 0.0022). **(G)** The CeL section of a *Prkcd-Cre;Ai14* mouse with *post hoc* immunostaining of SST and biocytin. Identified biocytin filled PKC-δ(+)/SST- and PKC-δ(-)/SST+ CeL neurons were depicted (#1 and #2, respectively). Scale bar, 20 μm. **(H)** Representative mIPSC recordings from PKC-δ(-) (red) and PKC-δ(+) (black) neurons in the CeL of Ctrl and FC-1d groups. **(I)** mIPSC frequency of CeL neurons of Ctrl and FC-1d *Prkcd-Cre;Ai14* mice (PKC-δ(+), Ctrl: 0.37 ± 0.08 Hz, n = 11 cells; FC-1d: 0.92 ± 0.16 Hz, n = 17 cells; PKC-δ(-), Ctrl: 0.35 ± 0.05 Hz, n = 11 cells; FC-1d: 0.39 ± 0.08 Hz; n = 13 cells; *p < 0.05, **p < 0.01, two-way ANOVA with Tukey’s multiple comparisons test). **(J)** mIPSC amplitude of CeL neurons of Ctrl and FC-1d groups *Prkcd-Cre;Ai14* mice (PKC-δ(+), Ctrl: 36.81.94 ± 4.22 pA, n = 11 cells; FC-1d: 44.56 ± 2.41 pA, n = 17 cells; PKC-δ(-), Ctrl: 3.03 ± 4.48 pA, n = 11 cells; FC-1d: 45.27 ± 2.62 pA; n = 13 cells; main treatment effect: Ctrl vs. FC, ^#^p = 0.0214, two-way ANOVA). **(K)** Plot of mIPSC frequency of PKC-δ(+) neurons against animal freezing levels (n = 10 animals, linear regression R square = 0.483, p = 0.026).

First, we crossed Sst-IRES-Cre mice with the Ai14 reporter mouse line to visualize SST+ CeL neurons by their tdTomato expression (Figure 2B). We also filled the recorded neurons with biocytin for *post hoc* identification (Figure 2B). We found that fear memory retrieval was accompanied by an increase in the frequency of mIPSCs recorded from SST-[mainly PKC-δ(+)], but not from SST+ [mostly PKC-δ(-)], CeL neurons at both 24 hrs (Figures 2C, D, S2A) and 7 days (Figures S1E-G) after FC. In contrast, FC elicited a general increment of mIPSC amplitude in both SST+ and SST-CeL neurons at both 24 hrs (Figure 2E, main treatment effect: Ctrl vs. FC, p = 0.0046, two-way ANOVA) and 7 days after FC (Figure S1H, main effect: Ctrl vs. FC, p = 0.0026, Two-way ANOVA). Similar results were observed when *ex vivo* recordings were performed 3 hrs after FC in animals that were not subjected to fear memory retrieval (Figures S1A-D), suggesting that inhibitory plasticity occurs shortly after the FC.

We also crossed Prkcd-Cre mice with the Ai14 reporter mouse line to directly access the PKC-δ(+) CeL neurons by their tdTomato expression (Figure 2G). In these mice, we observed that fear memory retrieval was accompanied by an increase in the mIPSC frequency in PKC-δ(+) (mainly SST-), but not in PKC-δ(-) (mostly SST+) CeL neurons recorded 24 hrs after FC (Figures 2H and 2I). On the other hand, FC elicited a general increment of the mIPSC amplitude in both PKC-δ(+) and PKC-δ(-) CeL neurons 24 hrs after FC (Figure 2J, main treatment effect: Ctrl vs. FC, p = 0.0214, two-way ANOVA).

Taken together, these results demonstrate that fear memory is accompanied by a preferential enhancement of mIPSC frequency onto CeL PKC-δ(+) neurons over SST+ neurons as well as a general postsynaptic increment of mIPSC amplitude of CeL neurons. Furthermore, we found that the mean mIPSC frequency of CeL SST- or PKC-δ(+) neurons can serve as an index to predict the degree of animal’s freezing during the fear memory retrieval test (Figures 2F and 2K). Since synaptic potentiation of excitatory synaptic transmission recorded from CeL SST+ and evoked by LA stimulation underlies fear memory (Li et al., 2013), we postulate that the inhibitory plasticity, including enhanced spontaneous IPSCs and mIPSCs, preferentially detected in PKC-δ(+) neurons also helps facilitate the biased activation towards CeL SST+ neurons after FC.

### Detection of fear memory CeL inhibitory engrams

Next, we advanced the hypothesis of the occurrence of an inhibitory engram in the CeL as a cellular substrate of fear memory. To test this hypothesis, we first examined whether FC increases CeL neuron activity by detecting the expression of the immediate early gene c-*fos*, a marker for recently activated neurons in various brain areas including the amygdala (Busti et al., 2011). We found that 12.75 ± 2.10% of CeL cells expressed c-Fos within 1.5 hour after FC, indicating a sparse labelling pattern. FC significantly increased the number of c-Fos+ cells in the CeL compared to the CS-only group (Cue; Figures S3A-C). This result represents an initial clue of the presence of an inhibitory engram in the CeL activated by fear memory.

Next, we sought to label and manipulate fear memory-activated neurons by using the targeted recombination in active populations method (TRAP2, DeNardo et al., 2019). First, we bilaterally injected in the CeL of TRAP2 mice an AAV-dlx-DIO-hKORD-cyRFP transducing these neurons to express an engineered kappa-opioid receptor that can suppress neuronal activity upon the administration of the designer receptors exclusively activated by designer drugs (DREADD) agonist salvinorin B (SALB; 10 mg/kg, s.c.), tagged with the red fluorescence protein (RFP). This viral vector used the dlx promoter to restrict the expression of hKORD-cyRFP only in CeL GABAergic neurons that displayed strong c-Fos activity induced by FC (FC-TRAPed cells).

Four weeks after injection, the experimental mice received a 4-hydroxytamoxifen (4-OHT) injection immediately after FC to induce Cre-dependent recombination (Figure 3A). Immunohistochemical analysis showed that the number of TRAPed cells was comparable to that of c-Fos+ cells in wild type mice one hour after FC (Figure 3C, Figure S3C). Furthermore, the number of CeL TRAPed cells in the FC-TRAP group was significantly higher than in CS-only controls (Cue-TRAP; Figures 3B and 3C).

**Figure 3.**
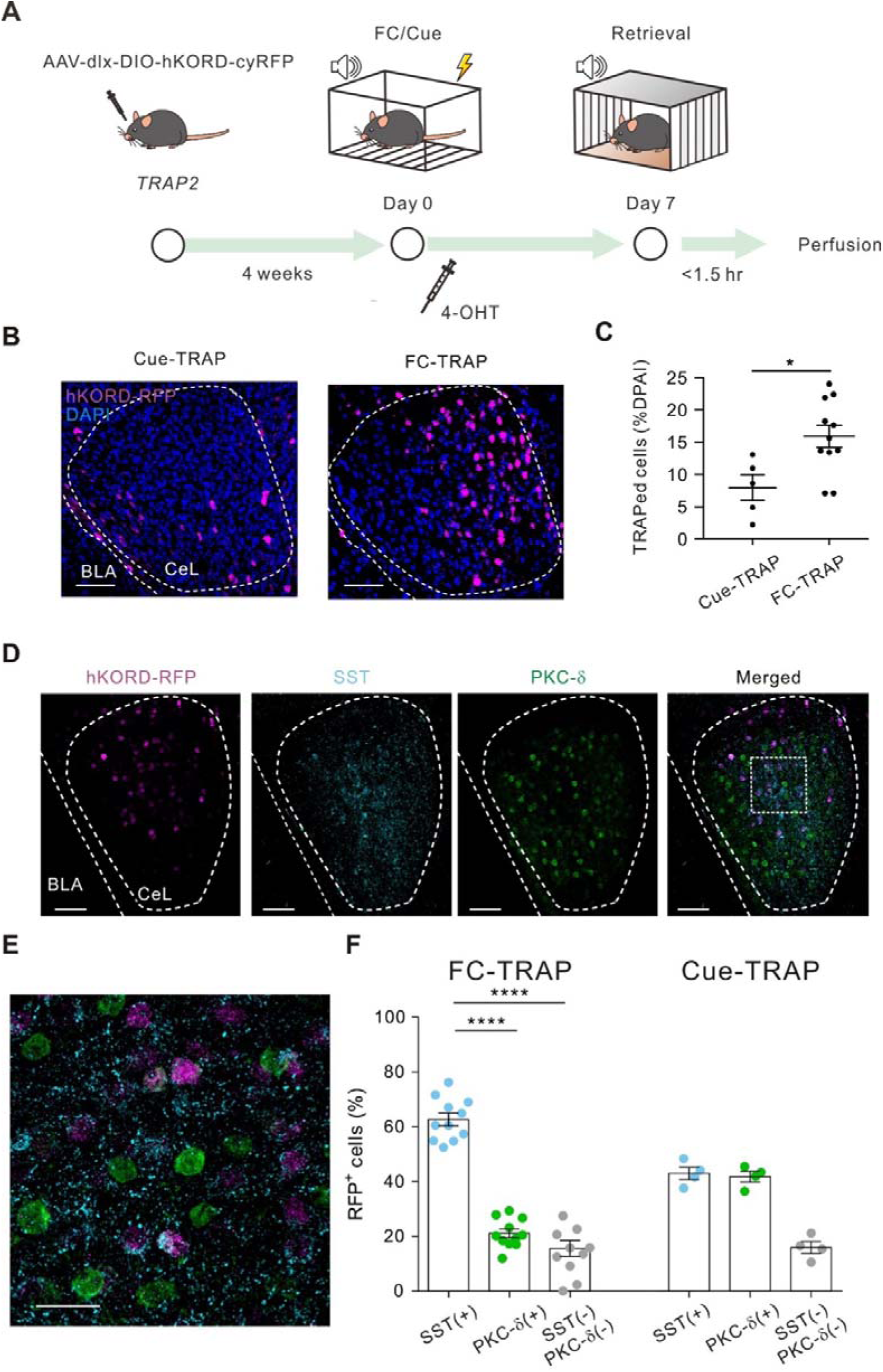
Tagging the CeL fear engram cells by TRAP2. **(A)** Experimental schematic and timeline. **(B)** Representative images of the TRAP pattern in the mouse CeL of Cue and FC groups. Scale bar, 50 μm. **(C)** Quantification of TRAPed cell proportion in the CeL (Cue-TRAP, 7.39 ± 1.58 % of total CeL cells, n = 5 mice, FC-TRAP, 15.76 ± 1.23 % of total CeL cells, n = 11 mice; *p < 0.05, Mann-Whitney test). **(D)** Characterization of the molecular identities of FC-TRAPed CeL neurons. Merged view of RFP (magenta), SST (cyan), and PKC-δ (green) using triple-immunostaining under 10X and **(E)** 63X magnification. Scale bars, (D) 100 μm; (E) 25 μm. **(F)** Quantification of the molecular identification of RFP+ TRAPed CeL neurons (FC-TRAP: SST+, 62.64 ± 2.32 %, PKC-δ(+), 21.05 ± 1.58 %, SST-/ PKC-δ(-), 15.55 ± 2.91 %; Cue-TRAP: SST+, 42.88 ± 2.24 %, PKC-δ(+), 41.73. ± 1.93 %, SST-/ PKC-δ(-), 15.90 ± 2.17 %; ****p < 0.0001, two-way ANOVA with Tukey’s multiple comparisons test).

Could the TRAPed cells represent the cellular substrate of a CeL inhibitory engram underlying fear memory retrieval 7 days after FC? Previous studies revealed that memory ensembles in cortical brain regions can be dynamic (De Sousa et al., 2019). Therefore, we tested whether the CeL engram remained stable or changed over time. To address this question, TRAP2 mice were subjected to FC with 4-OHT administration four weeks after the bilateral injection of AAV-dlx-DIO-hKORD-cyRFP in the CeL, and fear memory retrieval was tested 7 days later (Figure S3D). Mice were then perfused with a fixative and their brains processed for immunohistochemical analysis. This consisted of combined RFP and c-Fos immunostaining to determine the proportion of CeL TRAPed neurons activated by FC and reactivated by fear memory retrieval (Figure S3E). We found that FC-TRAPed(+) CeL neurons were more likely to express c-Fos upon memory retrieval compared to the FC-TRAPed(-) cells (Figures S3F-H), suggesting a certain stability of the CeL cells activated by FC. Interestingly, similar differences were observed in the Cue-TRAP group that showed a lower proportion of c-Fos immunolabeled cells compared to the FC-TRAP group, suggesting that a small fraction CeL neurons also encode information about the CS (Figures S3F-H).

Distinct experiences are known to be allocated into different engram populations in the neocortex, hippocampal dentate gyrus (DG), and amygdala (Josselyn and Tonegawa, 2020). We further asked if different CS-US associations recruited similar or distinct CeL cell clusters. The TRAP2 mice were bilaterally injected with AAV-dlx-DIO-hKORD-cyRFP and subjected to two different auditory cue–US pairings (CS1: 4 kHz and CS2: 12 kHz). We tagged the activated cells by the CS1-US pairing (FC1) with 4-OHT injection immediately after the FC1. After 7 days the mice were subjected to CS2-US pairing (FC2) and the c-Fos expression patterns were quantified as an indicator for FC2 activated neurons (Figure S2I). The mice showed comparable pre-conditioning freezing level and similar learning curves during FC1 and FC2 acquisition (Figure S3J). We next compared the FC1 activated (TRAPed) cells and the FC2 activated (c-Fos+) CeL cells. Despite the proportion of CeL TRAPed and c-Fos+ cells were comparable (TRAP: 13.14 ± 0.82 %, c-Fos: 15.53 ± 0.80 %), they were largely non-overlapping (TRAP+ c-Fos+: 2.28 ± 0.05 %, Figures S3K, L). Furthermore, the overlapping rate was close to the chance level (1.12 ± 0.03, n = 4 animals Figure S3M). These results indicate that the CeL is able to signal distinct CS-US associations and allocate them into largely non-overlapping neuronal ensembles.

### Neuron type composition of fear memory CeL inhibitory engrams

Since fear memory retrieval requires the activation of CeL SST+ neurons (Li et al., 2013) and induces synaptic plasticity in SST-/PKC-δ(+) neurons, we reasoned that the CeL engram should preferentially comprise SST+/PKC-δ(-) neurons. To test this possibility, we used the same fear memory paradigm shown in Figure S2D performed in TRAP2 mice bilaterally injected with AAV-dlx-DIO-hKORD-cyRFP into the CeL. Then, we examined the neurochemical features of CeL TRAPed neurons. Immunohistochemical analysis revealed that 62.64 ± 2.32 % of FC-TRAPed CeL cells (RFP+ cells) and 42.88 ± 2.24 % of the Cue-TRAP CeL cells were SST+ cells. In addition, 21.05 ± 1.58 % of CeL FC-TRAPed cells and 41.73 ± 1.93 % of the Cue-TRAP cells were PKC-δ(+). A smaller fraction of double negative cells was observed in both groups (15.55 ± 2.91 %, FC-TRAP; 15.90 ± 2.17%, Cue-TRAP; Figure 3D-F). Statistical comparison revealed that FC preferentially activated CeL SST+ neurons (Chi-square test, Figure 3F). It is worth noting that the TRAPed SST+ CeL cells only accounted for 18% of total SST+ cells. In conclusion, CeL FC-TRAPed cells, though sparse and heterogeneous, comprise preferentially, but not exclusively, SST+ neurons.

### Anatomical output of CeL GABAergic fear engram neurons

We then investigated the areas that are predominantly targeted by CeL FC-TRAPed neurons. As mentioned previously, CeL SST+ neurons represent the majority of engram cells, and have been shown to project both locally within the CeL (Li et al., 2013; Hou et al., 2016; Hunt et al., 2017) and to extra-amygdaloid areas (Penzo et al., 2014; Sun et al., 2020b; Figure 4A). Therefore, we tested whether a similar pattern of axonal projection would be represented by CeL fear engram neurons. To examine CeL FC-TRAPed neuron projection patterns, we utilized the expression of hKORD-cyRFP by CeL FC-TRAPed neurons (Figure 4B). We detected dense axonal branches of SST+ neurons (Figure 4C and Figure S4C) and CeL FC-TRAPed neurons within the CeL (Figure S4D), suggesting that these terminals could represent a major source of GABA release to elicit mIPSCs recorded from CeL neurons in our electrophysiological experiments. Our findings also revealed CeL FC-TRAPed neuron efferents in several extra-amygdaloid areas, that are also targeted by axon terminals of CeL SST+ neurons. They included: the bed nucleus of stria terminalis (BNST), the periaqueductal gray (PAG), the lateral habenula (LHb) and the dorsal medial geniculate nucleus (MGd, Figure 4D-G, right panels) and several posterior thalamic nuclei (PoT, PIL, Po and PF, Figure 4G, right panel; Figure S4A). In addition, we observed labelling in the nuclei of the horizontal and vertical limb of the diagonal band (HDB and VDB; Figure 4 G, H, right panels; Figure 4J) and the arcuate nucleus (ARH, Figure S4B, right panel). Overall, our results suggest that the projection pattern of FC engram cells followed that of CeL SST+ neurons (Figure 4D-H, left panels; Figure 4I), consistent with the finding that the majority of CeL engram cells are SST+.

**Figure 4.**
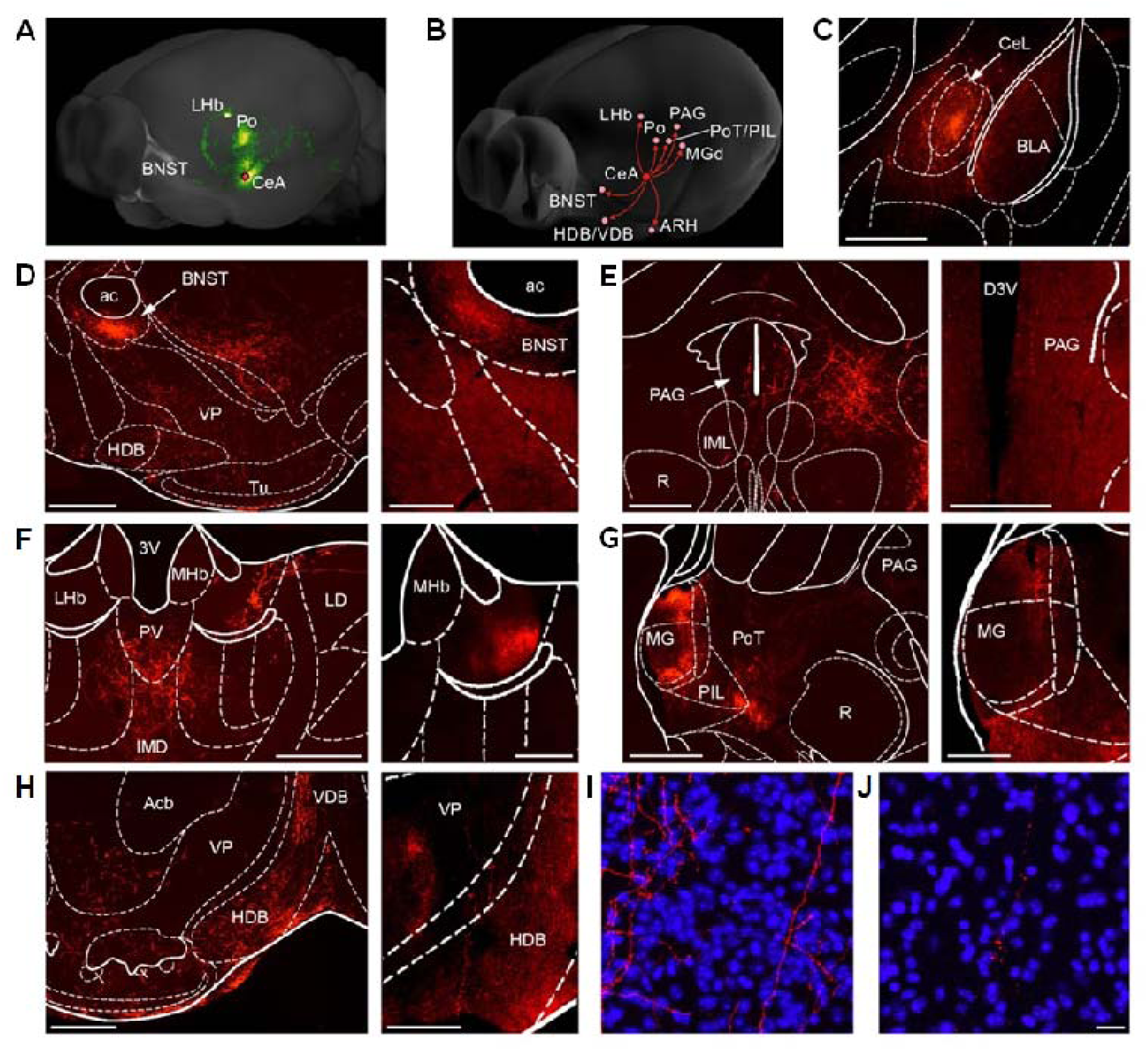
Whole brain mapping of axonal projections from CeL inhibitory engram neurons. **(A)** 3D view of CeL SST+ neuron projections obtained from the Allen Mouse Brain Connectivity Atlas (experiment 539641136). Image credit: Allen Institute. The pattern of innervation of transduced CeL SST+ neurons was highly similar to the one observed in our tracing experiments using the viral vector AAV-DIO-hChR2-mCherry. **(B)** 3D overview of brain areas receiving inputs from CeL FC-TRAPed neurons, as illustrated here and in the Supplemental Figure 4. **(C)** Micrograph of the injection site of the viral vector AAV-DIO-hChR2-mCherry in the CeL from a representative Sst-IRES-Cre mouse. Scale bar, 500 μm. **(D-H)** Examples of brain areas that were consistently found targeted by hChR2-mCherry-labeled axons in Sst-IRES-Cre mice (panels on the left) as well as by hKORD-RFP-labeled axons from FC-TRAPed mice (panels on the right). Scale bars, (D-H) (left) 500 μm; (D-G) (right) 250 μm; (H) (right) 200 μm. **(D)** Low magnification images displaying a high density of labeled axon terminals from CeL SST+ neurons (left, 5x) or from FC-TRAPed neurons in the lateroventral subdivision of the BNST (right). **(E)** Labeled axons coming from CeL SST+ neurons (left) or FC-TRAPed neurons (right) can also be detected in the PAG. **(F)** A dense innervation from SST+ (left) and TRAPed neurons (right) appears confined to the lateral LHb. **(G)** The posterior intralaminar thalamic nuclei (PoT and PIL) and subareas of the MG (dorsal part) receive numerous labeled axons from CeL SST+ neurons (left). Likewise, the same areas are targeted by FC-TRAPed neurons (right). **(H)** A number of labeled axonal branches from CeL SST+ neurons can be seen running through the nuclei of the HDB and VDB (left). Axons of FC-TRAPed neurons are also observed in these nuclei (right). **(I-J)** Confocal micrographs displaying representative examples of axonal branches and terminals in sections from the HDB counterstained with DAPI (in blue) in Sst-IRES-Cre (I) and FC-TRAPed (J) mice. Thickness of confocal z stacks: (I) 39.4 μm; (J) 17.8 μm. Scale bar, 20 μm. Abbreviations: ac, anterior commissure; Acb, nucleus accumbens; BLA, basolateral amygdaloid nucleus; BNST, bed nucleus of stria terminalis; CeA, central amygdala; CeL, central amygdaloid nucleus lateral part; HDB, nucleus of the horizontal limb of the diagonal band; IMD, intermediodorsal thalamic nucleus; IML, interstitial nucleus of the medial longitudinal fasciculus; LD, laterodorsal thalamic nucleus; LHb, lateral habenular nucleus; MHb, medial habenular nucleus; MG, medial geniculate nucleus; PAG, periaqueductal grey area; PIL, posterior intralaminar thalamic nucleus; Po, posterior thalamic nuclear group; PoT, post-thalamic nucleus; R, red nucleus; PV, paraventricular thalamic nucleus; Tu, olfactory tubercle; VDB, nucleus of the vertical limb of the diagonal band; VP, ventral pallidum.

### Silencing the CeL inhibitory fear engram increases mouse freezing

Given that CeL FC-TRAPed cells are activated upon fear memory retrieval, our next objective was to examine the impact of silencing the inhibitory engram on fear memory. To this end, we microinjected TRAP2 mice with AAV-dlx-DIO-hKORD-cyRFP in the CeL and treated them systemically with 4-OHT immediately after FC to label the CeL FC engram We then divided the mice into two experimental groups: one group received an injection of 10 mg/kg SALB, while the other served as the control group and received a vehicle injection (Veh). Both groups were injected either with SALB or Veh 30 minutes before fear memory retrieval, which occurred 7 days after FC (Figure 5A). Our *in vitro* electrophysiological experiments revealed that SALB-induced hKORD activation led to membrane hyperpolarization and reduced firing in CeL neurons recorded in acute slices (Figure S5A-F). During fear acquisition, both groups of mice displayed similar levels of freezing (Figure 5B). However, the silencing of CeL FC-TRAPed cells through SALB administration during fear memory retrieval resulted in a significant increase in freezing compared to both the Veh-treated FC-TRAPed group and the SALB group exposed only to the CS (Figure 5C). Notably, there were no significant differences in pre-CS baseline freezing levels between the experimental groups before memory retrieval at 7 days after FC (Figure 5C). Further control experiments were performed to rule out that chemogenetic inactivation of CeL FC-TRAPed cells induced generalized spontaneous freezing or altered motor activity in the open field test (OFT, Figures S5G-J).

**Figure 5.**
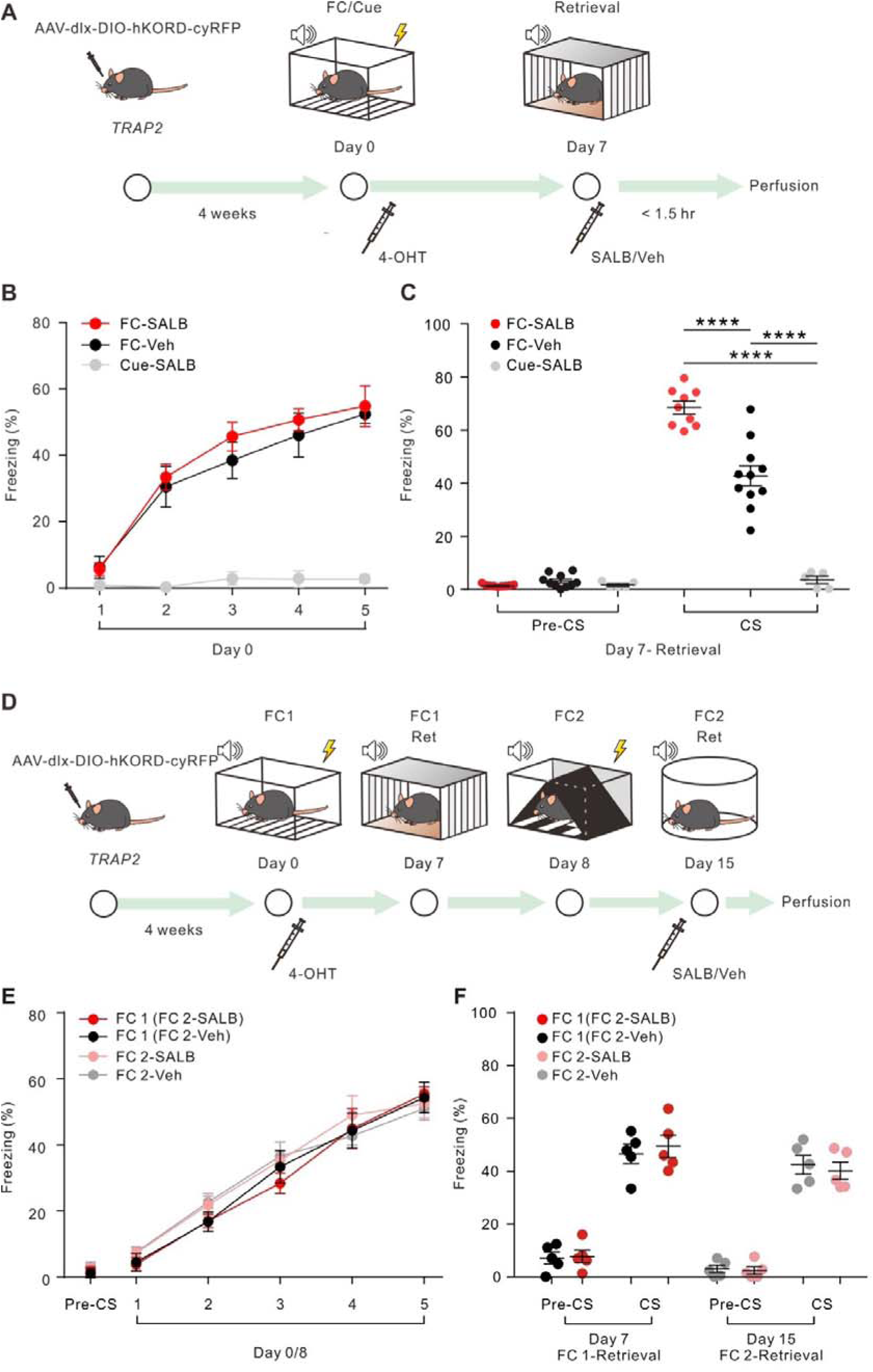
Chemogenetic suppression of CeL inhibitory engram enhanced CS-dependent freezing behavior. **(A)** Experimental schematic and timeline. **(B)** Freezing (time in %) during fear conditioning. FC-SALB, n = 9 animals; FC-Veh, n = 11 animals; Cue-SALB, n = 5 animals); p < 0.0001, two-way ANOVA. **(C)** Freezing (time in %) during memory retrieval on day 7. Pre-CS: FC-SALB, 1.80 ± 2.41 %; FC-Veh, 3.10 ± 4.85 %; Cue-SALB, 0.80 ± 0.50 %; CS-presentation: FC-SALB, 68.51 ± 2.35 %; FC-Veh, 42.83 ± 3.81 %; Cue-SALB, 3.36 ± 1.41 %; two-way ANOVA with Tukey’s multiple comparisons test. ****p < 0.0001. **(D)** Experimental schematic and timeline. **(E)** Freezing (time in %) during fear conditioning. FC 1 (FC2-SALB) group, n = 5 animals; FC1 (FC2-Veh) group, n = 5 animals; FC 2-SALB, n = 5 animals; FC2-Veh, n = 5 animals. **(F)** Freezing (time in %) during memory retrieval on day 7. Pre-CS: FC1 (FC2-SALB), 7.74 ± 2.39 %; FC1 (FC2-Veh), 7.13 ± 2.25 %; FC2-SALB, 2.40 ± 1.42 %; FC2-Veh, 3.02 ± 1.38 %. CS-presentation: FC1 (FC2-SALB), 51.99 ± 4.24 %; FC1 (FC2-Veh), 49.09 ± 3.65 %; FC2-SALB, 40.21 ± 3.23 %; FC2-Veh, 42.88 ± 3.54 %; two-way ANOVA with Tukey’s multiple comparison test.

We further tested the specificity of the effects induced by silencing the inhibitory engram by using the two CS-US association paradigms described previously (Figure 5D). We chemogenetically inactivated the FC1 CeL engram during the retrieval of another memory with a similar behavioral outcome (FC2, Figure 5D). The mice were transfected with AAV-dlx-DIO-hKORD-cyRFP and underwent FC1 and subsequent 4-OHT injection and memory retrieval on day 7 as shown in Figure 5D. On the next day, the mice were subjected to FC2. Seven days after FC2, the mice were split into Veh-and SALB-treated groups for the FC2 retrieval. Both groups exhibited similar freezing level during the acquisition of FC1 and FC2, and retrieval of FC1 (Figure 5E, F). During the FC2 retrieval, both Veh- and SALB-treated groups showed low and similar freezing to the testing context for FC2 (Pre-CS, Figure 5F). Moreover, Veh- and SALB-treated groups froze to a similar degree to CS2 presentations (CS, Figure 5F). In an additional set of experiments, in which mice that underwent the same two CS-US association paradigm (Figure S6A) and received SALB injections 30 minutes prior to both the FC1 and FC2 retrieval, we could confirm the specificity of the increased freezing response upon the FC1 (similar to the result shown in Figure 5C), but not the FC2 retrieval (Figures S6A-C). Furthermore, the SALB injection in the TRAP2 mice without hKORD expression did not alter fear learning and freezing level upon fear memory retrieval (Figures S7A-C). Compared to the hKORD expression groups (Figures S5G-J), SALB injection did not alter the anxiety level and motor activity in the non-hKORD expression TRAP2 mice (Figures S7D-F). Taken together, these findings let us to conclude that the enhanced freezing level induced by silencing the CeL inhibitory engram is specific to a distinctive CS-US association.

Finally, we also investigated whether there is an engram-specific change in mIPSC frequency in the engram population. We measured mIPSCs from engram and non-engram CeL neuron pairs in the same slice using *TRAP2;Ai14* mice (7 cell pairs from 7 slice, 5 mice, Figures S8A-C). Our results revealed that the mIPSC frequency of TRAP(-) cells was significantly higher in TRAP(-) cells than in TRAP(+) cells. Therefore, our findings suggest that the change in mIPSC frequency occurs mainly in non-engram cells.

### Suppression of CeL FC-TRAPed neurons disinhibits downstream target regions

Considering that CeL FC-TRAPed cells are GABAergic, we reasoned that, in addition to local inhibitory synaptic plasticity within the CeL, they could influence the activity of downstream brain regions, for example by disinhibition. Indeed, we observed that the number of c-Fos+ cells was significantly higher compared to that of the Veh-treated group in several areas targeted by the CeL FC-TRAPed neuron efferents. These included the local CeL, ventral BNST (vBNST), PAG, LHb, MGd, MGv, and ARH (Figure 6A-G). Thus, CeL FC-TRAPed neurons are likely to depress downstream extra-amygdaloid inhibitory neurons resulting in disinhibition of downstream target neurons.

**Figure 6.**
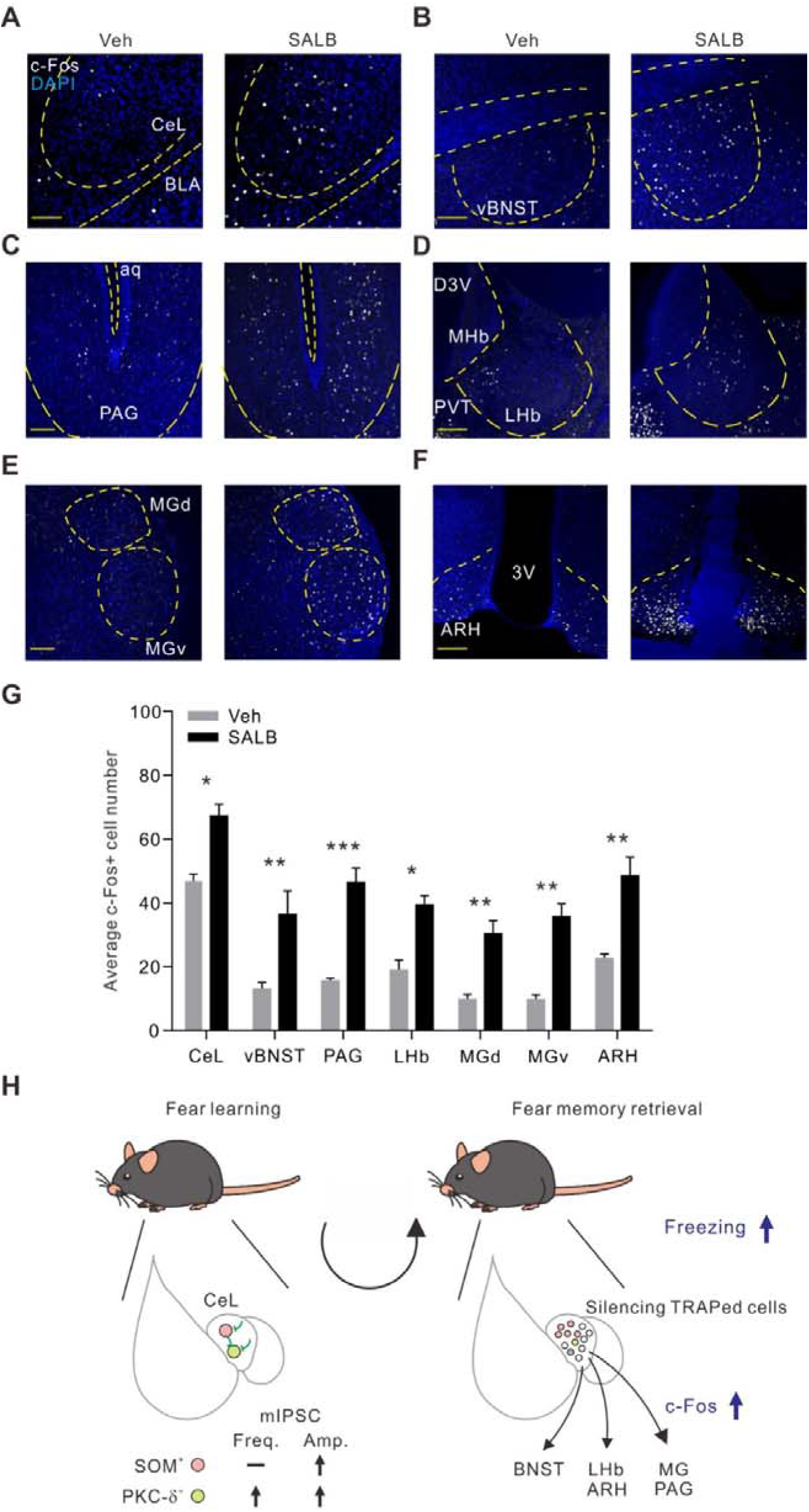
Chemogenetic silencing CeL FC-TRAPed neurons enhanced the c-Fos expression in the target areas. **(A-F)** Representative images of c-Fos expression in the target regions of the CeL FC-TRAPed GABAergic cells by either Veh or SALB injection 30 mins before 7-day fear memory test. (A), CeL; (B), ventral BNST (vBNST); (C), PAG; (D), LHb; (E), dorsal and ventral MG (MGd and MGv); (F), ARH. Scale bars, 100 μm. **(G)** Quantification of average c-Fos+ cell numbers: CeL: Veh 46.91 ± 2.13 cells, SALB 63.98 ± 5.35 cells, p < 0.05; vBNST: Veh, 13.00 ± 2.08 cells, SALB, 36.42 ± 7.34 cells, p < 0.01; PAG: Veh, 15.50 ± 0.88 cells, SALB, 46.42 ± 4.49 cells, p < 0.001; LHb: Veh, 18.92 ± 3.11 cells, SALB, 39.42 ± 2.87 cells, p < 0.05; MGd: Veh, 9.75 ± 1.56 cells, SALB, 30.33 ± 4.12 cells, p < 0.05; MGv: Veh, 9.72 ± 1.48 cells, SALB, 35.83 ± 3.98 cells, p < 0.01; ARH: Veh, 22.67 ± 1.33 cells, SALB, 48.42 ± 5.85 cells, p < 0.01; two-way ANOVA with Tukey’s multiple comparison test. n = 3 animals. *p < 0.05; **p < 0.01; ***p < 0.001. **(H)** Working hypothesis of CeL inhibitory fear engram. Left, fear conditioning enhanced the activity of a subset CeL neurons that are mainly SST+ (pink). The sustained elevated activity of those cells (inhibitory engram) is necessary for fear memory formation, and is likely to contribute to long-term GABAergic plasticity, leading to increased net inhibition onto CeL PKC-δ(+) cells (light green) and extra-amygdaloid target neurons. Right, silencing TRAPed cells (inhibitory engram) causes disinhibition of extra-amygdaloid target neurons and increases in their c-Fos numbers and freezing levels. Abbreviations: ARH, arcuate region of the hypothalamus; BNST, bed nucleus of stria terminalis; LHb, lateral habenular nucleus; MG, medial geniculate nucleus; PAG, periaqueductal grey area.

Collectively, our findings provide compelling evidence for the existence of a fear memory engram within the CeL region, primarily comprised of SST+ neurons (as shown in Figure 6H). We further demonstrate that this engram is characterized by enhanced inhibitory synaptic transmission that targets both local CeL PKC-δ(+) neurons and extra-amygdaloid neurons, ultimately leading to the suppression of downstream circuits and the inhibition of fear memory expression.

## DISCUSSION

Our study has identified a unique fear memory engram formed exclusively by GABAergic neurons within the CeL region of mice. We demonstrate that the temporary suppression of this inhibitory engram results in increased disinhibition of targeted areas, including extra-amygdaloid regions, ultimately leading to the facilitation of fear memory expression, as evidenced by an increase in freezing behavior in mice.

The central amygdala (CeA) is a key area for the acquisition, consolidation, and expression of fear memories (Ciocchi et al., 2010; Duvarci and Pare, 2014; Herry et al., 2008; Tovote et al., 2015). The current model proposes that after FC, the CS is processed by the BLA (Sah et al., 2003; Tovote et al., 2015), but the conditioned response is mediated by excitatory connections to SST+ neurons of the CeL (Li et al., 2013) that in turn inhibit local PKC-δ(+) neurons (Ciocchi et al., 2010). According to this model, the inhibitory projections from the PKC-δ(+) neurons disinhibit CeM neurons (Ciocchi et al., 2010; Haubensak et al., 2010), but also project to the LA (Yu et al., 2017), to elicit defensive responses through connections to downstream targets.

Our data are consistent with this model and further add novel aspects to it. Firstly, we demonstrate that after FC, only a subset of CeL neurons (∼13% of total CeL cells) are activated. Their activity is not only present upon fear memory retrieval 24 hrs later (Ciocchi et al., 2010), but also at 3 hrs and even at 7 days after FC. Secondly, our results show that FC induces long-lasting inhibitory synaptic plasticity, mainly onto SST-/PKC-δ (+) CeL neurons. Thirdly, fear memory retrieval preferentially activates SST+ neurons, despite the fact that the percentage of GABAergic cells expressing SST or PKC-δ appears to be highly similar in the mouse CeL (Shrestha et al., 2020). Fourthly, we demonstrate that inhibition of CeL engram neurons enhances fear expression. Lastly, our results indicate that SST+ neurons not only inhibit local neurons, but also send widespread connections to extra-amygdaloid areas that serve as relay nodes of the distributed freezing pathway (Penzo et al., 2014; Tovote et al., 2016).

Our findings are consistent with a recent report demonstrating monosynaptic inhibitory projections from CeL SST+ neurons to neurons of the central sublenticular extended amygdala (Sun et al., 2020b). Notably, most of the extra-amygdaloid areas targeted by CeL FC-TRAPed neurons are also critical sites for generalization of auditory fear memory, such as CeM, MG, and PIL (Asok et al., 2019; Barsy et al., 2020). It is also important to note that the projection patterns of the CeL FC-TRAPed are distinct from the CeA ensemble activated by general anesthesia, that was primarily composed of PKC-δ(+) neurons, as previously reported (Hua et al., 2020).

We show that chemogenetic inhibition of CeL FC engram neurons results in increased freezing behavior during fear memory retrieval 7 days after FC. This finding supports the idea that engrams formed by inhibitory neurons may actually “mask” memory (Barron et al., 2017). Downstream areas targeted by the CeL FC-TRAPed neurons, such as the LHb (Lecca et al., 2017), PIL (Lanuza et al., 2004), and PAG (Tovote et al., 2016), have previously been shown to be activated by footshocks and that their activity is required for freezing behaviour. In keeping with this, our c-Fos data suggest that silencing of CeL FC-TRAPed neurons promotes the disinhibition of downstream regions, leading to increased freezing behavior during fear memory retrieval. We also observed elevated c-Fos activity in several thalamic nuclei, including the posterior intralaminar thalamic nuclei (PoT, in addition to PIL) and the MGd, when the CeL FC-TRAPed neurons were chemogenetically silenced. These connections may close a loop that involves thalamic nuclei and the LA, extending to the CeL to integrate the association of CS and US signals (Barsy et al., 2020). In addition to extra-amygdaloid mechanisms, disinhibition within the CeL itself may also contribute to the observed increase in freezing behavior. CeL SST+ neurons are highly interconnected (Hou et al.,2016; Hunt et al.,2017), and chemogenetic silencing of a small fraction of these neurons may result in the disinhibition of other SST+ neurons, leading to overall increased freezing. Furthermore, the disinhibition of CeM output neurons by PKC-δ(+) neurons, which are part of the CeL FC inhibitory engram, may represent another contributing mechanism. Moreover, we did not observe enhanced pre-CS freezing during retrieval test, suggesting that silencing of the CeL FC engram does not trigger context-dependent fear generalization (Figure 5F).

Our work is one of the few documented cases of inhibitory synaptic plasticity following an associative learning paradigm, such as FC. A previous study have demonstrated long-term potentiation (LTP) of cortical PV-expressing basket cell-mediated synaptic inhibition in response to visual deprivation (Maffei et al., 2006). Furthermore, recent research has shown that cortical spike timing dependent plasticity of PV+ interneuron-mediated inhibition contributes to auditory map remodeling (Vickers et al., 2018). Similarly, prefrontal SST+ interneurons have been reported to encode fear memory (Cummings and Clem, 2020; Cummings et al, 2022). Although we did not thoroughly investigate the mechanisms underlying fear memory-mediated long-term inhibitory synaptic plasticity, our results indicate that the frequency of mIPSCs were potentiated by fear memory, predominantly in PKC-δ(+) neurons. This points to a potential target-specific presynaptic mechanism (Thompson et al., 1993). Concurrently, FC also induces a non-specific mIPSC amplitude enhancement in CeL neurons, which may act as a general adaptation to the heightened plasticity of glutamatergic synapses following fear learning. This could be accompanied by a possible parallel increase in GABA synapses and/or GABA-A receptors. In relation to this, research has demonstrated that FC triggers an increase in the proportion of synaptic benzodiazepine-sensitive GABA-A receptors with the α2 subunit in pyramidal neurons of the basal amygdala (Kasugai et al., 2019). As a result, future studies should assess whether this molecular mechanism, or another, is functional in the CeL after FC.

Our findings identify an inhibitory engram activated by fear memory retrieval, predominantly formed by SST+ neurons of the CeL. Using the TRAP2 approach, we found that FC leads to a proportion of TRAPed neurons similar to that of c-Fos+ cells in wild type mice (∼13% of total CeL cells), which is consistent with a recent study (DeNardo et al., 2019). Thus, the CeL inhibitory engram involved in fear memory retrieval appears sparse, in line with the small size of engrams formed by excitatory neurons (Rao-Ruiz et al., 2019). These include 2–6% of neurons in the dentate gyrus (Liu et al., 2012; Tayler et al., 2013) and 10–20% in the hippocampal CA1 region (Tayler et al., 2013), LA (Gouty-Colomer et al., 2016; Han et al., 2007) or prefrontal cortex (Kitamura et al., 2017) as cellular substrates of conditioned fear memory. The sparse feature of the inhibitory CeL engram may explain some conflicting results in the literature. For example, PKC-δ(+) CeL neurons have been proposed to be either “fear-off” (Ciocchi et al., 2010; Haubensak et al., 2010), namely displaying inhibitory CS responses following FC and acting to suppress fear responses (Ciocchi et al., 2010; Haubensak et al., 2010) or “fear-on” (Yu et al., 2017), namely delivering aversive US signals, driving aversive learning, activated by the CS and predicting the US (Yu et al., 2017). Our results suggest that only a small percentage of PKC-δ(+) CeL neurons is recruited by an inhibitory engram to encode a specific fear memory. This makes the idea of the co-existence of non-overlapping “fear-on” and “fear-off” PKC-δ(+) CeL neurons less problematic.

The role of inhibitory engrams is currently debated (Josselyn and Tonegawa, 2020). So far, engrams formed by inhibitory neurons have been mostly studied in circuits expressing both glutamatergic and GABAergic cells (Cummings and Clem, 2020; Cummings et al, 2022). In this context, GABAergic engram neurons have been proposed to play a role in keeping the balance or controlling the extension of glutamatergic excitatory engrams (Barron et al., 2017; Josselyn and Tonegawa, 2020; Rao-Ruiz et al., 2019). For example, the size of a LA fear excitatory engram is regulated by local PV(+) interneurons during FC (Morrison et al., 2016). Furthermore, the inhibition of SST+ interneurons during contextual FC increases the size of a hippocampal excitatory dentate granule cells engram (Stefanelli et al., 2016). Consequently, modeling studies have proposed that inhibitory engrams play a role in keeping the excitatory-inhibitory balance during memory processes (Barron et al., 2017). However, it is possible that inhibitory engrams can also have distinct functional roles, as compared to excitatory engrams. Furthermore, cortical inhibitory engrams may prevent unwanted co-activation of overlapping memories (Koolschijn et al., 2019). Computational evidence suggested that GABAergic engrams might serve an opposite role to excitatory engrams and inhibit memory (Barron et al., 2017).

A key question is understanding how neurons are selected to become part of an engram (Rao-Ruiz et al., 2019). This does not only have considerable theoretical relevance, it also informs computational models, and it may facilitate the artificial manipulation of engrams for future therapeutic interventions. Both experimental and computational data suggest that neuron’s intrinsic excitability and synaptic strength are the most critical factors in promoting engram selection (Bocchio et al., 2017; Poo et al., 2016; Rao-Ruiz et al., 2019). For example, LA neurons with high intrinsic excitability (Yiu et al., 2014) or those connected with strong excitatory synaptic interactions are more likely to be incorporated into an engram (Kim et al., 2013). This latter result is in line with engram formation rules demonstrated in hippocampal granule cells (Ryan et al., 2015). Our results suggest that synaptic inhibition also plays a role in engram formation. Neurons receive less inhibition, e.g. CeL SST+ neurons, are more likely to contribute to an engram. Another potential factor influencing the engram formation is the strength of the inputs. It is currently unclear which inputs activate CeL inhibitory engram neurons. Previous studies suggested that FC enhances excitatory synapses onto SST+ neurons and weakens those onto SST− neurons in the CeL (Li et al., 2013). Moreover, the PVT innervation of CeL SST+ neurons plays a crucial role in the expression of fear learning (Penzo et al., 2015).

Finally, it is important to acknowledge some methodological limitations of our study. The labeling of behaviorally activated neurons by TRAP2 may be dependent on the brain area, due to varying c-Fos activity levels resulting in high or low signal thresholds. Additionally, the type of activity that triggers the c-Fos signal might be relatively undefined, potentially encompassing plastic synaptic events as well as burst and tonic neuronal firing (Kawashima et al., 2014). Despite these limitations, our study’s detection of TRAPed cells in the mouse CeL after FC aligns well with previous findings (DeNardo et al., 2019). Another limitation is that our *in vitro* experiments were conducted shortly after a high fear state. Under these conditions, we cannot rule out the possibility that the observed synaptic changes may reflect alterations in the physiological/behavioral state associated with fear expression (Goosens et al.,2003). Nevertheless, we believe that our identification of an inhibitory engram in the mouse CeL of the amygdala underlying fear memory significantly contribute to understanding the role of engrams formed by GABAergic neurons. This novel observation warrants further investigation. For example, the inhibitory engram could be studied using an in *situ* multi-electrode array recording approach designed to selectively capture the activity of engaged neurons.

## ACKNOWLEDGEMENTS

This work was supported by grants to M.C. (AUFF NOVA 2016; PROMEMO – Center for Proteins in Memory, a Center of Excellence funded by the Danish National Research Foundation grant number DNRF133) to F.F. (Austrian Science Fund grant number I-2215) to C.C.L. (supported by the Brain Research Center, National Yang Ming Chiao Tung University from The Featured Areas Research Center Program within the framework of the Higher Education Sprout Project by the Ministry of Education in Taiwan, National Health Research Institutes (NHRI-EX110-10814NI), and Ministry of Science and Technology (108-2320-B-010 -026 -MY3; MOST 110-2321-B-010-006) in Taiwan) and to W.H.H. (Ministry of Science and Technology, Taiwan, fellowship 110-2917-I-564-011). We acknowledge the technical support by Kathrine Meinecke Christensen, Majken Sand, and Stella Solveig Nolte. We also thank Emma Louth, Meike Sieburg (Capogna’s lab), and Sadegh Nabavi (DANDRITE Aarhus) for their comments on the manuscript. The authors report no conflict of interest.

## AUTHOR CONTRIBUTION

Conceptualization, W.H.H., C.C.L., and M.C.; Investigation, W.H.H., M.J., K.Y.W., A.S., Y.L.L. and A.R.; Formal analysis and data visualization, W.H.H., M.J., K.Y.W., A.S., Y.L.L. and A.R., Writing-original draft, W.H.H. and M.C.; Writing-review & editing, W.H.H., M.J., F.F., C.C.L. and M.C.; Funding acquisition, W.H.H., C.C.L., F.F. and M.C.; Supervision, C.C.L., F.F. and M.C..

## DECLARATION OF INTERESTS

The authors declare no competing interests.

## Supplemental information

**Figure S1.**
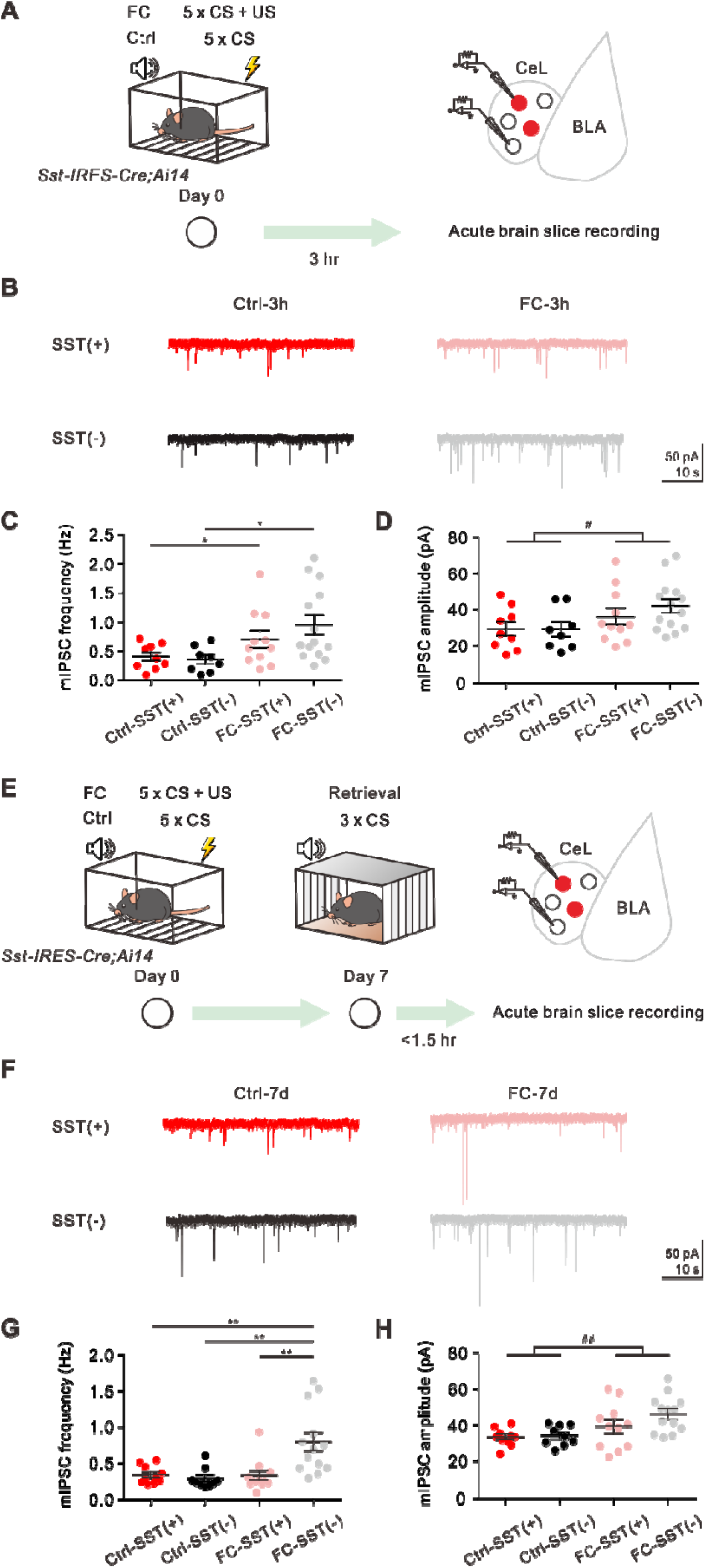
Enhanced mIPSCs 3 hrs and 7 days after fear conditioning. Related to Figure 2. **(A)** Experimental schematic and timeline. **(B)** Representative mIPSC recordings from SST+ (red traces) and SST− (black traces) neurons in the CeL of Ctrl and FC-3h groups. **(C)** mIPSC frequency of CeL neurons of Ctrl and FC-3h groups (SST+, Ctrl: 0.39 ± 0.07 Hz, n = 9 cells, 4 animals; FC-3h: 0.69 ± 0.15 Hz, n = 11 cells, 6 animals; SST−, Ctrl: 0.34 ± 0.08 Hz, n = 8 cells, 4 animals; FC-3h: 0.93 ± 0.17 Hz; n = 14 cells, 6 animals; *p < 0.05, two-way ANOVA with Tukey’s multiple comparisons test). **(D)** mIPSC amplitude of CeL neurons of Ctrl and FC-3h groups (SST+, Ctrl: 37.81 ± 4.76 pA, n = 9 cells, 4 animals; FC-3h: 46.03 ± 5.51 pA, n = 11 cells, 6 animals; SOM−, Ctrl: 37.49 ± 5.03 pA, n = 8 cells, 4 animals; FC-3h: 53.3 ± 4.61 pA; n = 14 cells, 6 animals; main treatment effect: Ctrl vs. FC, ^#^p = 0.026, two-way ANOVA). **(E)** Experimental schematic and timeline. **(F)** Representative mIPSC recordings from SST+ (red traces) and SST− (black traces) neurons in the CeL of Ctrl and FC-7d groups. **(G)** mIPSC frequency of CeL neurons of Ctrl and FC-7d groups (SST+, Ctrl: 0.3322 ± 0.04 Hz, n = 10 cells, 2 animals; FC-7d: 0.32 ± 0.07 Hz, n = 11 cells, 2 animals; SST−, Ctrl: 0.28 ± 0.05 Hz, n = 9 cells, 2 animals; FC-7d: 0.80 ± 0.13 Hz; n = 13 cells, 2 animals; **p < 0.01, two-way ANOVA with Tukey’s multiple comparisons test). **(H)** mIPSC amplitude of CeL neurons of Ctrl and FC-7d groups (SST+, Ctrl: 31.57 ± 1.52 pA, n = 10 cells, 2 animals; FC-7d: 37.58 ± 3.78 pA, n = 11 cells, 2 animals; SST−, Ctrl: 29.29 ± 1.86 pA, n = 9 cells, 2 animals; FC-7d: 44.56 ± 2.80 pA; n = 13 cells, 2 animals; main treatment effect: Ctrl vs. FC, ^##^p = 0.0026, two-way ANOVA).

**Figure S2.**
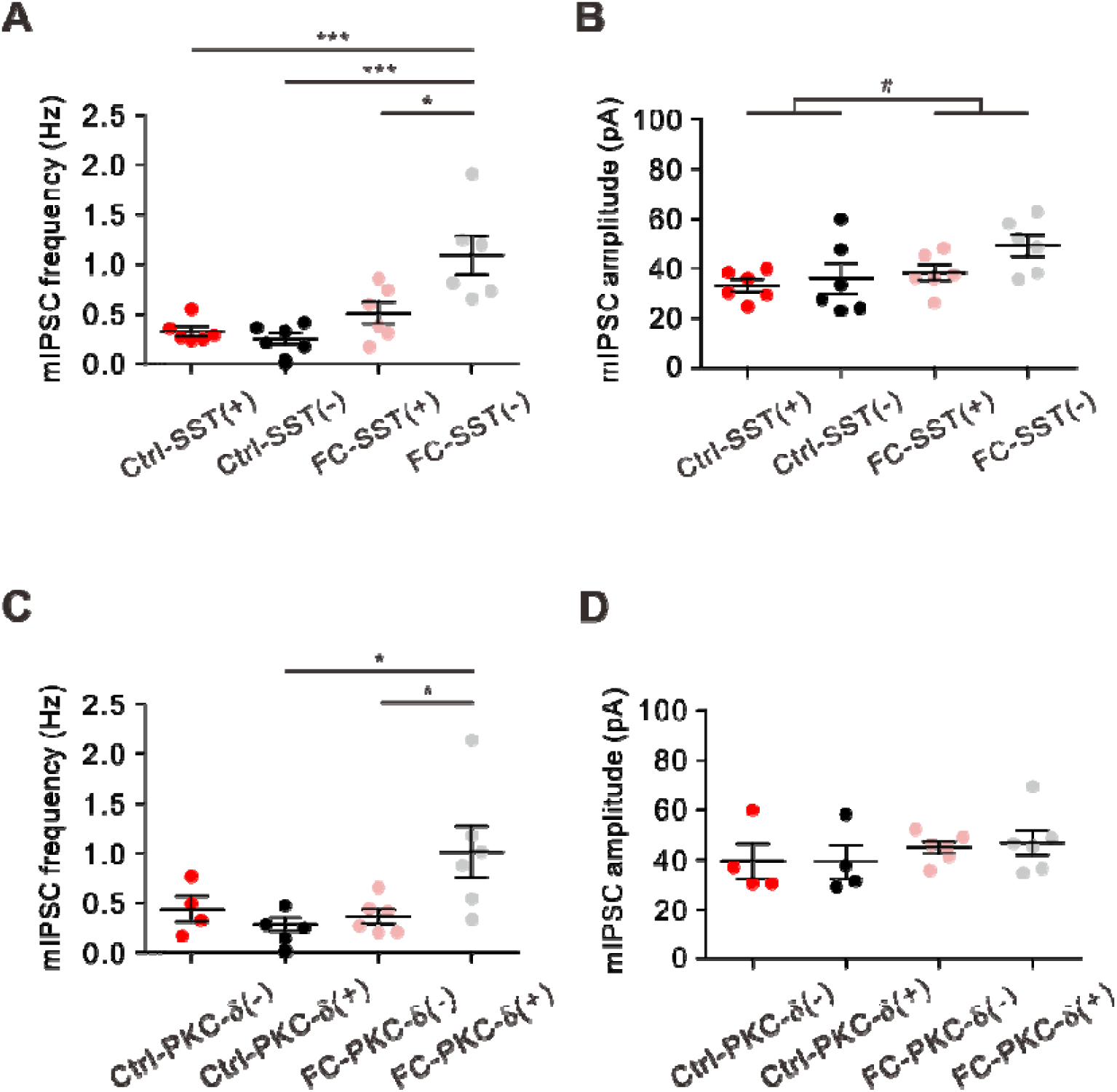
mIPSC data in Fig. 2 presented by animal average. Related to Figure 2. **(A)** mIPSC frequency of CeL neurons of Ctrl and FC-1d groups (SST+, Ctrl: 0.33 ± 0.049 Hz, n = 6 animals; FC-1d: 0.51 ± 0.11 Hz, n = 6 animals; SST−, Ctrl: 0.26 ± 0.06 Hz, n = 6 animals; FC-1d: 1.09 ± 0.19 Hz, n = 6 animals; *p < 0.05, ***p < 0.001, two-way ANOVA with Tukey’s multiple comparisons test). **(B)** mIPSC amplitude of CeL neurons of Ctrl and FC-1d groups (SST+, Ctrl: 33.22 ± 2.43 pA, n = 6 animals; FC-1d: 36.04 ± 6.02 pA, n = 6 animals; SST−, Ctrl: 49.34 ± 4.37 pA, n = 6 animals; FC-1d: 52.03 ± 5.56 pA, n = 6 animals; main treatment effect: Ctrl vs. FC, ^#^p = 0.0409, two-way ANOVA). **(C)** mIPSC frequency of CeL neurons of Ctrl and FC-1d Prkcd-Cre;Ai14 mice (PKC-δ(+), Ctrl: 0.29 ± 0.07 Hz, n = 4 animals ; FC-1d: 1.01 ± 0.26 Hz, n = 6 animals; PKC-δ(-), Ctrl: 0.44 ± 0.13 Hz, n = 4 animals ; FC-1d: 0.37 ± 0.07 Hz; n = 6 animals; *p < 0.05, two-way ANOVA with Tukey’s multiple comparisons test). **(D)** mIPSC amplitude of CeL neurons of Ctrl and FC-1d groups Prkcd-Cre;Ai14 mice (PKC-δ(+), Ctrl: 39.10 ± 6.62 pA, n = 4 animals ; FC-1d: 46.80 ± 5.07 pA, n = 6 animals ; PKC-δ(-), Ctrl: 39.48 ± 7.00 pA, n = 4 animals ; FC-1d: 44.79 ± 2.39 pA; n = 6 animals; two-way ANOVA with Tukey’s multiple comparisons test).

**Figure S3.**
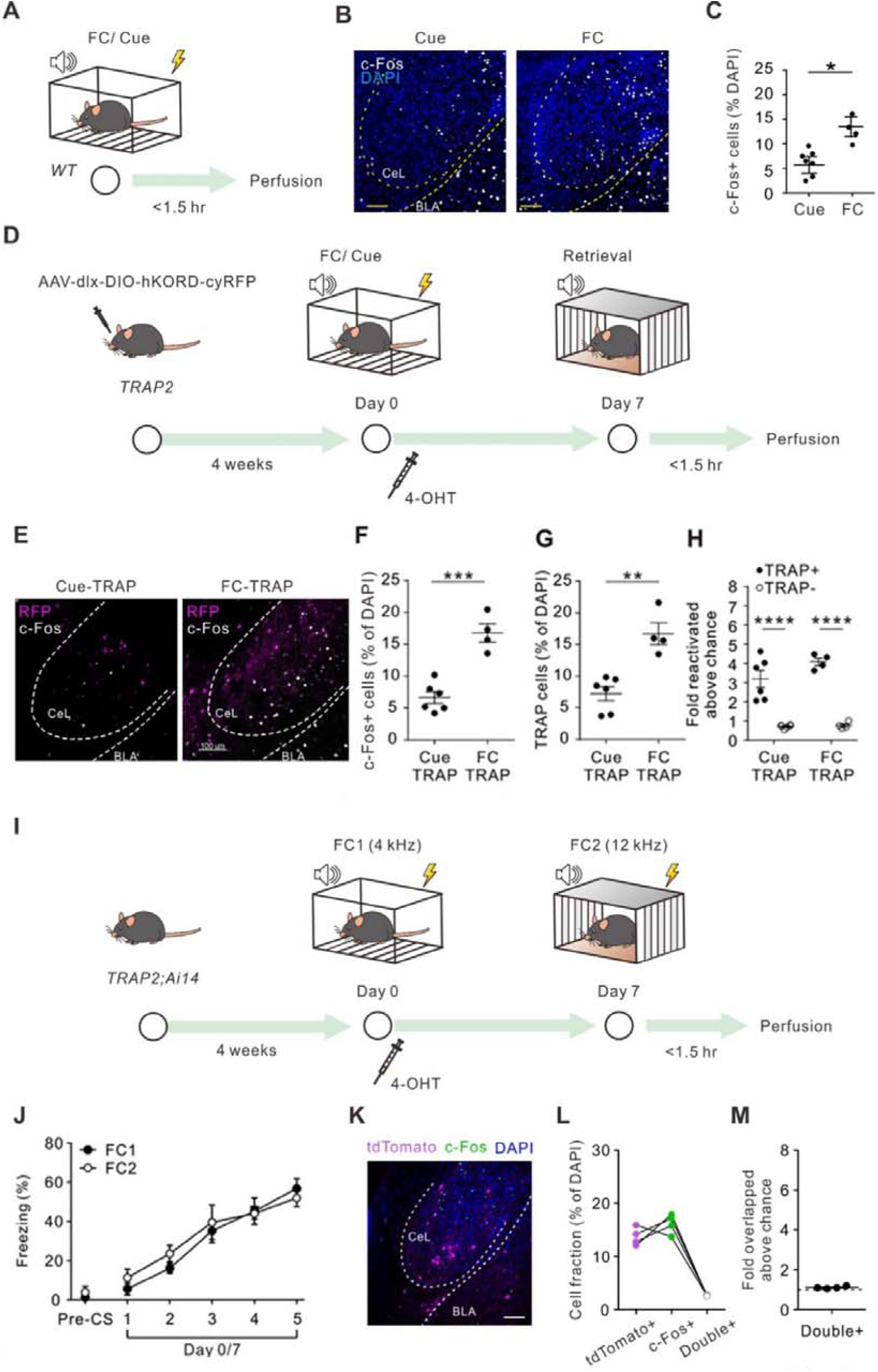
Activation of Fos+ and TRAP2+ cells after fear conditioning. Related to Figure 3. **(A)** Fear conditioning paradigm. **(B)** Representative images of c-Fos expression of cells from both Cue and FC groups 90 min after FC. Scale bar, 50 μm. **(C)** The proportion of c-Fos+ cells in both Cue and FC groups was quantified (Cue: 5.93 ± 1.07 % of total CeL cells, n = 7 mice, FC: 12.75 ± 2.10 % of total CeL cells, n = 4 mice; p = 0.02, Mann-Whitney test. *p < 0.05. **(D)** Fear conditioning TRAP experimental paradigm. **(E)** Representative images of TRAPed (RFP+) and c-Fos+ cells from both Cue-TRAP and FC-TRAP groups. Scale bar, 100 μm. **(F)** Reactivation rate of CeL FC-TRAPed cells during the retrieval of 7-day fear memory was quantified (Cue-TRAP, 36.53 ± 3.96 % of total CeL cells, n = 6 mice, FC-TRAP, 64.90 ± 4.21 % of total CeL cells, n = 4 mice; p = 0.0095, Mann-Whitney test). **p < 0.01. **(G)** Co-localization fraction of FC-TRAPed cells with the c-Fos+ cells in CeL was quantified after memory retrieval on Day 7 (Cue-TRAP, 34.04 ± 2.81 % of total CeL cells, n = 6 mice, FC-TRAP, 64.82 ± 5.05 % of total CeL cells, n = 4 mice; p = 0.0095, Mann-Whitney test). **p < 0.01. **(H)** Comparison of c-Fos expression normalized to chance levels in TRAPed ((Double+/DAPI)/((c-Fos+/DAPI)x(TRAP+/DAPI)) and non-TRAPed ((Double+/DAPI)/((c-Fos+/DAPI)x(TRAP–/DAPI))) cells (two-way ANOVA with Tukey’s multiple comparisons test), **p < 0.01. **(I)** Experimental paradigm for distinct associative fear conditioning. On day 0, *TRAP2;Ai14* mice were subjected to auditory fear conditioning FC1 (5 x CS1 + US, CS1: 4 kHz) and 4-OHT injection. On Day 7, they were subjected to FC2 (5 x CS2 + US, CS2: 12 kHz) and were perfused within 90 minutes after the FC2 acquisition. **(J)** Average freezing time during pre-conditioning habituation and auditory associative fear conditioning. **(K)** Representative images of FC1-TRAPed (tdTomato+) and c-Fos+ cells. Scale bar, 50 μm. **(L)** Fraction of FC1-TRAPed (tdTomato+), c-Fos+, and co-localized cells in the mouse CeL. **(M)** The likelihood of colocalization normalized to chance levels.

**Figure S4.**
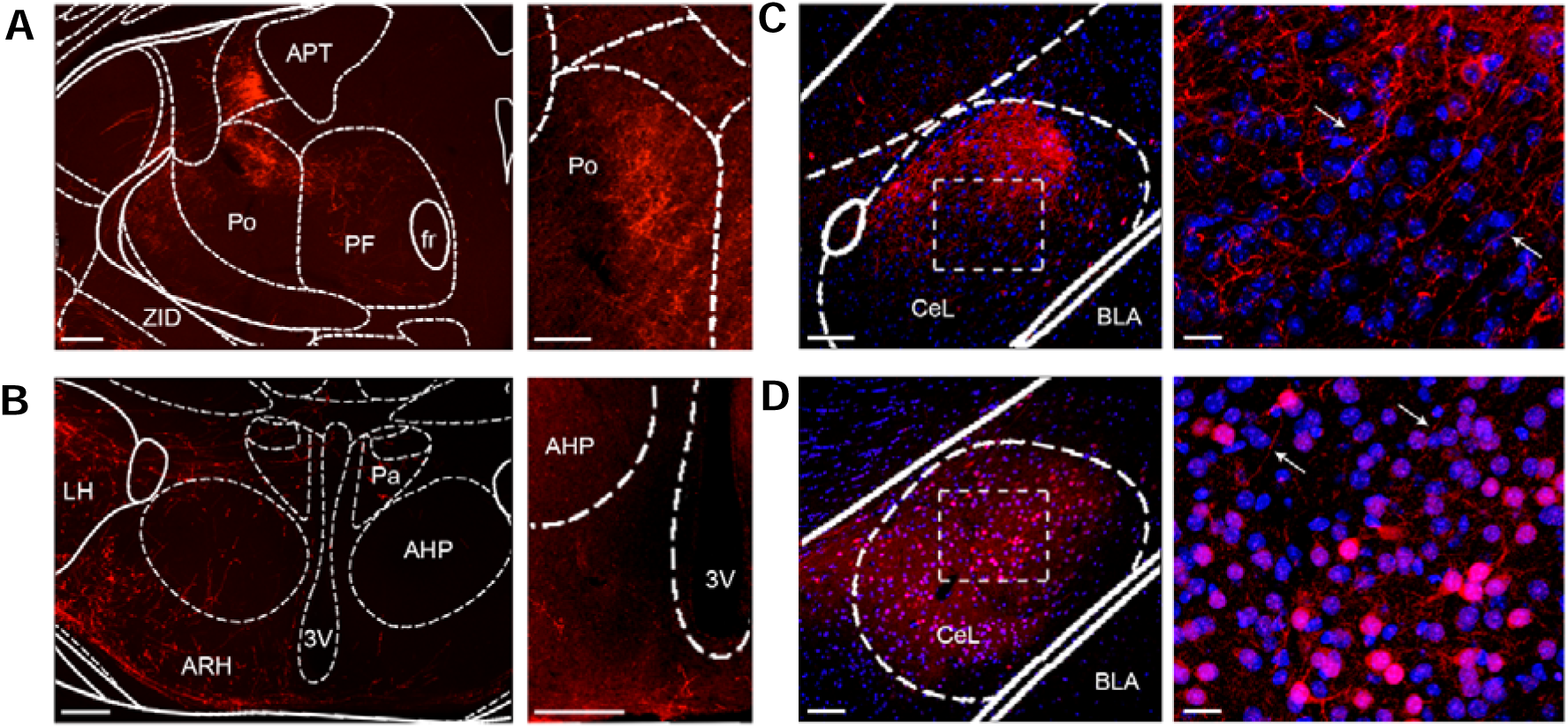
Further mapping of axonal projections from CeL inhibitory engram neurons. Related to Figure 4. **(A)** Labeled axonal branches originated from CeL SST+ neurons (left) can be observed in the Po, PF and the medio-rostral part of the lateral posterior thalamic nucleus. Axons of FC TRAPed neurons are particularly dense in the Po (right). **(B)** Sparse axonal labeling from CeL SST+ neurons (left) or FC TRAPed neurons (right) can also be detected in the ARH. Scale bars, (A-B) (left) 250 μm; (A) (right) 100 μm; (B) (right) 200 μm. **(C)** Labeling from CeL SST+ neurons (left) with zoomed-in local axonal distributions indicated by arrows (right). **(D)** Labeling from CeL TRAP+ neurons (left) with zoomed-in local axonal distributions indicated by arrows (right). Scale bars, (C-D) (left) 100 μm; (C-D) (right) 20 μm. Abbreviations: 3V, third ventricle, AHP, anterior hypothalamic area posterior part; APT, anterior pretectal nucleus; ARH, arcuate region of the hypothalamus; BLA, basolateral amygdala; CeL, central lateral amygdala; fr, fasciculus retroflexus; LH, lateral hypothalamic area; Pa, paraventricular hypothalamic nucleus; PF, parafascicular thalamic nucleus; Po, posterior thalamic nuclear group; ZID, zona incerta dorsal part.

**Figure S5.**
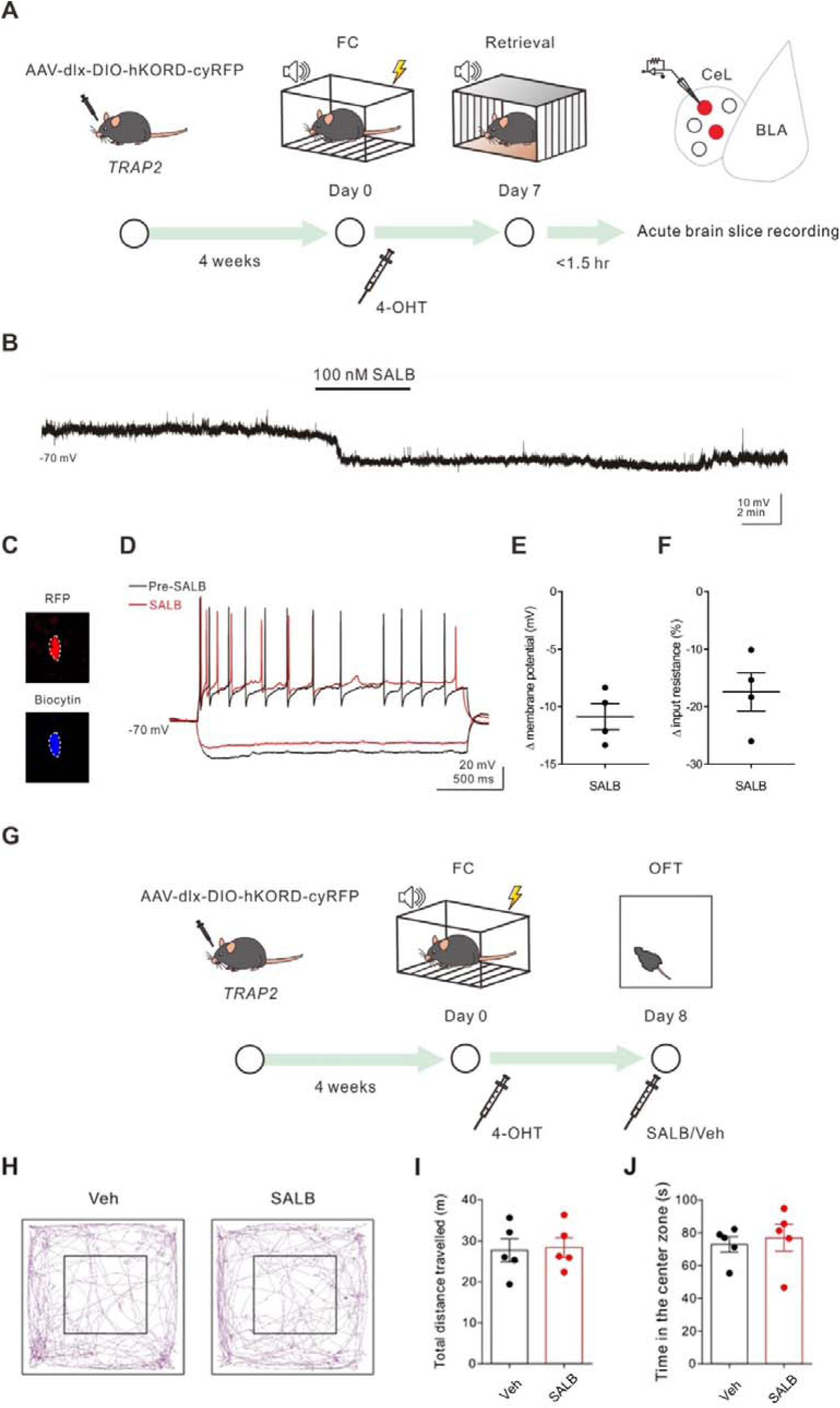
Effects of hKORD activation on the membrane potential of TRAP neurons *in vitro* and spontaneous locomotor activity *in vivo*. Related to Figure 5. **(A)** Experimental schematic and timeline. **(B)** Membrane potential changes of a hKORD-expressing CeL neuron recorded in the presence of kynurenic acid (2 mM) and gabazine (1 μM) in response to bath-applied SALB (100 nM). **(C)** Confocal images of a biocytin-filled RFP+ cells. Scale bar, 10 μm. **(D)** Firing patterns of the recorded cell before (black trace) and after (red trace) 100 nM SALB bath application. Note the decrease in input resistance after SALB application. **(E)** Net membrane potential change and **(F)** net input resistance change of the hKORD-expressing cells after the bath application of SALB (-10.87 ± 1.13 mV; - 17.46 ± 3.32%; n = 4 cells, 2 animals). **(G)** Experimental schematic and timeline. **(H)** Representative trajectories of Veh- and SALB-treated mice in the open field test. **(I)** Total travelled distance (Veh: 27.70 ± 2.83 m, n = 5 animals; SALB: 28.4 ± 2.43 m, n = 5 animals, p = 0.80, Mann-Whitney test) and **(J)** time in center zone (Veh: 72.92 ± 7.45 s, n = 5 animals; SALB: 76.98 ± 8.17 s, n = 5 animals, p = 0.68, Mann-Whitney test).

**Figure S6.**
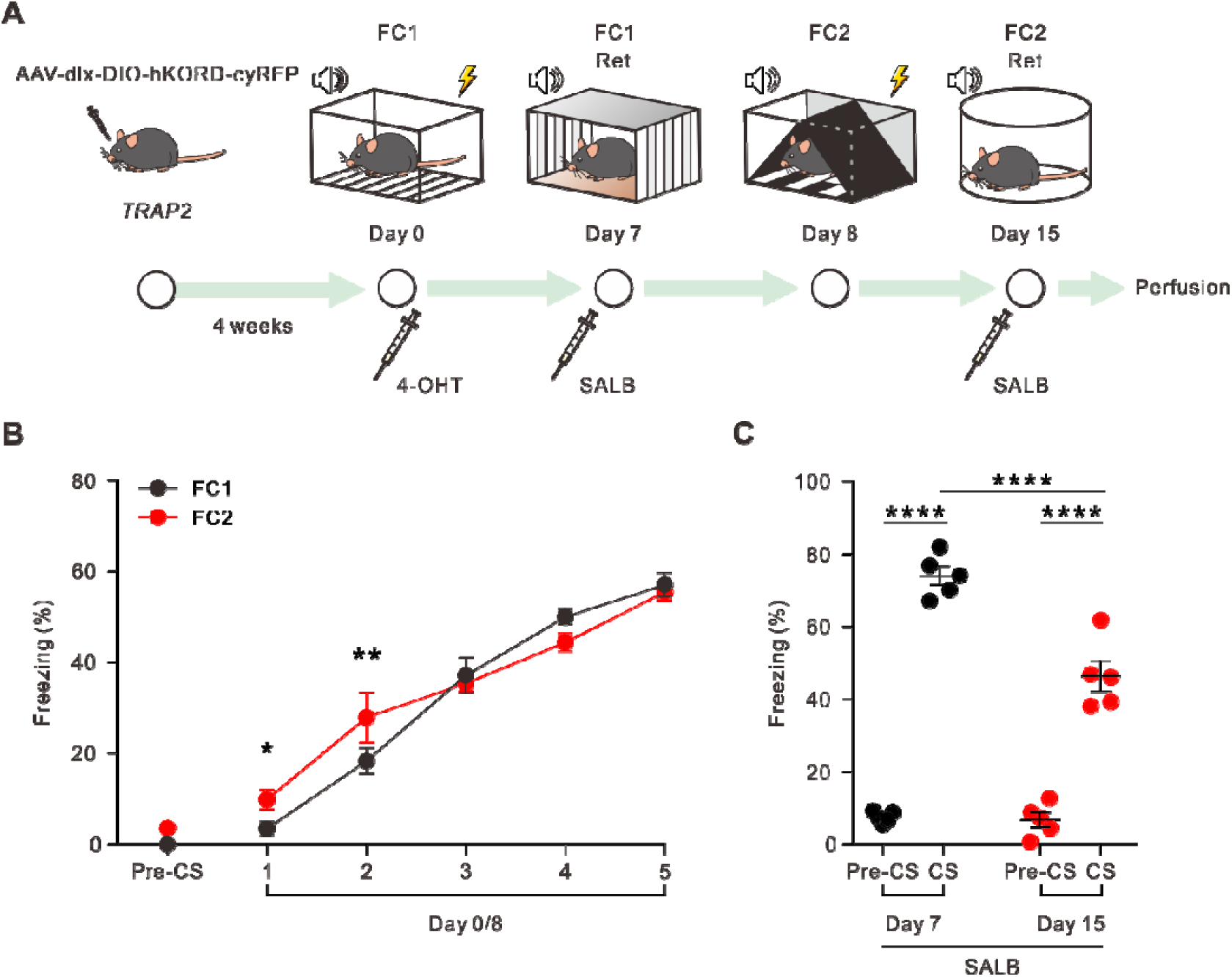
Chemogenetic suppression of CeL inhibitory engram enhanced CS-dependent freezing behavior without affecting distinct fear memory recall. Related to Figure 5. **(A)** Experimental schematic and timeline. **(B)** Freezing (time in %) during FC1 and FC2, n = 5 animals; two-way ANOVA with Tukey’s multiple comparison test. *p < 0.05, **p < 0.01. **(C)** Freezing (time in %) during memory retrieval on Day 7. Pre-CS: FC1-SALB, 7.60 ± 0.69 %; FC2-SALB, 6.82 ± 2.01 %. CS-presentation: FC1-SALB, 74.05 ± 2.59 %; FC2-SALB, 47.45 ± 4.23; two-way ANOVA with Tukey’s multiple comparison test. ****p < 0.0001.

**Figure S7.**
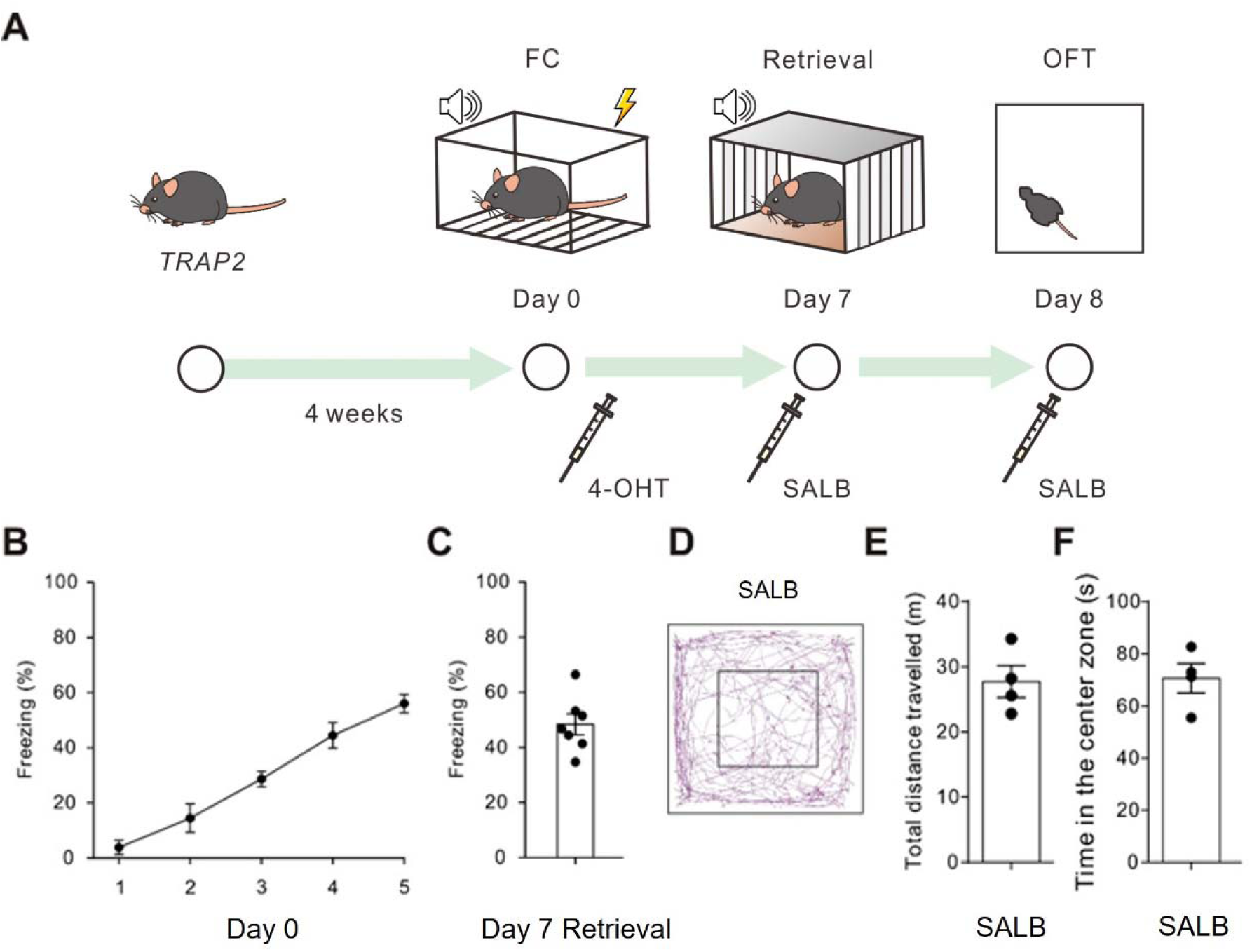
Further behavior examinations of SALB injection in the non-hKORD expressing TRAP2 mice. Related to Figure 5. **(A)** Experimental schematic and timeline. **(B)** Average freezing time during FC and **(C)** Memory retrieval on day 7 (n = 7 animals). **(D)** Representative trajectory, **(E)** Total travelled distance (27.6 ± 2.36 m), and **(F)** time in center zone (70.68 ± 5.13 s) of SALB-treated non-hKORD expressing mice in the open field test (n = 4 animals).

**Figure S8.**
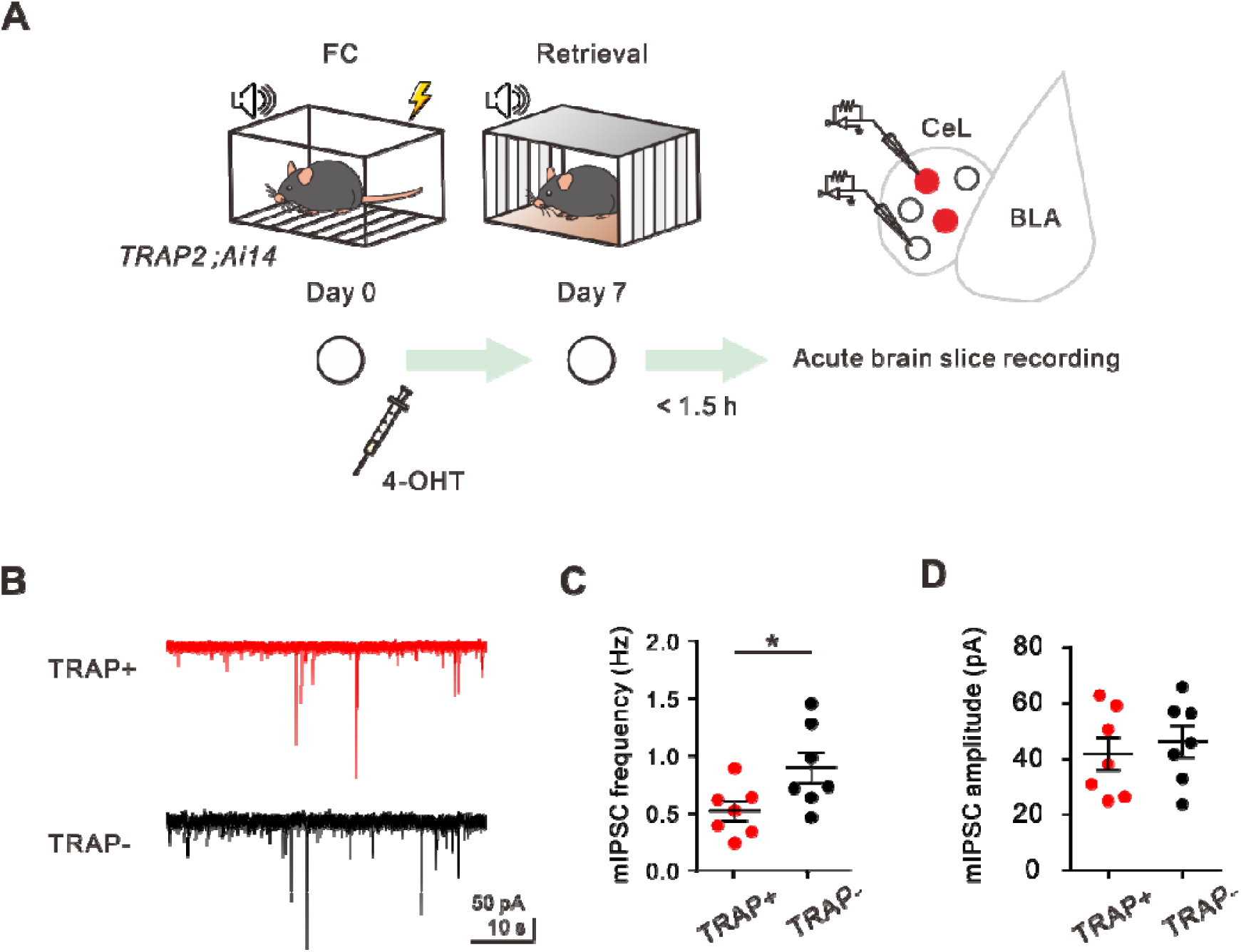
CeL FC-TRAP(+) cells showed less mIPSC frequency compared to TRAP(-) cells. Related to Figure 5. **(A)** Experimental schematic and timeline. **(B)** Representative mIPSC recordings from FC-TRAP(+) (red traces) and FC-TRAP(−) (black traces) neurons in the CeL. **(C)** Comparison of the mIPSC frequency between FC-TRAP(+) and FC-TRAP(-) neurons in the CeL. CeL FC-TRAP(-) cells show a significantly higher mIPSC frequency compared to FC-TRAP(+) cells (TRAP+: 0.52 ± 0.08 Hz, n = 7 cells; TRAP-: 0.90 ± 0.13 Hz, n = 7 cells, 5 animals; p = 0.0156, Wilcoxon matched-pairs signed rank test). *p < 0.05. **(D)** Comparison of the mIPSC amplitude between FC-TRAP(+) and FC-TRAP(-) neurons in the CeL. No significant difference in the mIPSC amplitude was observed between CeL FC-TRAP(+) and FC-TRAP(-) cells (TRAP+: 41.86 ± 5.92 pA, n = 7 cells; TRAP-: 46.22 ± 5.59 pA, n = 7 cells, 5 animals; p = 0.69, Wilcoxon matched-pairs signed rank test).

## Notes

### Competing Interest Statement

The authors have declared no competing interest.

