## Supplemental information for "Inhibitory fear memory engram in the mouse central lateral amygdala"

**Key Resources Table**

| Reagent or Resource | Source | Identifier |
|---|---|---|
| **Antibodies** | | |
| Rat monoclonal anti-Somatostatin (clone YC7) | Sigma-Aldrich | Cat# MAB354 |
| Mouse monoclonal anti- PKC-δ (clone 14) | BD Biosciences | Cat# 610397 |
| Rabbit polyclonal anti-c-Fos | Abcam | Cat# Ab190289 |
| Goat polyclonal anti-RFP | ORIGENE | Cat# AB1140-100 |
| Donkey anti-Goat Alexa Fluor 568 | Thermofisher | Cat# A-11057 |
| Donkey anti-Rabbit Cy5 | Jackson ImmunoResearch | Code: 711-175-152 |
| Donkey anti-Rat Alexa Fluor 488 | Sigma-Aldrich | Cat# SAB4600037 |
| Donkey anti-Mouse Alexa Fluor 488 | Jackson ImmunoResearch | Code: 715-545-150 |
| Donkey anti-Mouse Cy5 | Jackson ImmunoResearch | Code: 715-175-150 |
| Streptavidin-conjugated Alexa Fluor 488 | Thermofisher | Cat# S11223 |
| Streptavidin-conjugated Alexa Fluor 594 | Thermofisher | Cat# S11227 |
| **Bacterial and Virus Strains** | | |
| ssAAV-1/2-hDlx-HBB-chI-dlox-HA_hKORD_mCyRFP1(rev)-dlox-WPRE-hGHp(A) | VVF Zurich, Switzerland | v326-1 |
| ssAAV-5/2-mDlx-HBB-chI-dlox-hChR2(H134R)-mCherry(rev)-dlox-WPRE-bGHp(A) | VVF Zurich, Switzerland | v317-5 |
| **Chemicals, Peptides, and Recombinant Proteins** | | |
| Dimethyl sulfoxide | J.T.Baker | Cat# 9224-06 |
| Kynurenic acid | Sigma-Aldrich | Cat# K3375 |

| | | |
|---|---|---|
| Paraformaldehyde | Sigma-Aldrich | Cat# 158127 |
| Tetrodotoxin | Tocris | Cat# 1078 |
| Tetrodotoxin citrate | Abcam | Cat# ab120055 |
| Biocytin | Thermofisher | Cat# B-1592 |
| Salvinorin B (SALB) | Hello bio | Cat# HB4887 |
| Fentanyl (0.05 mg/kg) | Hameln | Cat# 007007 |
| Midazolam (5 mg/kg) | Hameln | Cat# 002124 |
| Medetomidine (0.5 mg/kg) | VM Pharma | Cat# 087896 |
| Naloxon (1.2 mg/kg) | B. Braun | Cat# 115241 |
| Atipamezole Hydrochloride (2.5 mg/kg) | Orion Pharma Animal Health | N/A |
| Flumazenil (0.5 mg/kg) | Sintetica | Cat# 4448335 |
| Xylocaine | AstraZeneca AB | Cat# 39699/0086 |
| Viscotears | Alcon Danmark A/S | Cat# 464110 |
| Buprenorphine (0.3 mg/ml) | Indivior | N/A |
| 4-Hydroxytamoxifen (4-OHT) | Sigma-Aldrich | Cat# H6278 |
| Mouse-on-Mouse blocking reagent (M.o.M) | Vector Laboratories | Cat# MKB 2213 |
| Normal Horse Serum | Sigma-Aldrich | Cat# H0146 |
| Normal Goat Serum | Vector Laboratories | Cat# S-1000 |
| Triton X-100 | USB Co. | N/A |
| VECTASHIELD Antifade Mounting Medium with DAPI | Vector Laboratories | Cat# H-1200 |
| Fluoromount-G™ Mounting Medium | Thermofisher | Cat# 00-4958-02 |
| **Experimental Models: Organisms/Strains** | | |
| Mouse: C57BL/6J | Janvier labs | N/A |
| Mouse: *Sst-IRES-Cre* | The Jackson Laboratory | Stock #013044 |
| Mouse: *Fos2A-iCreERT2* | The Jackson | Stock #030323 |

| | | |
|---|---|---|
| (TRAP2) | Laboratory | |
| Mouse: *Ai14* | The Jackson Laboratory | Stock #007908 |
| Mouse: *PKC:GluCl -ires-Cre* | Mutant Mouse Research and Resource Center | Stock #011559-UCD |
| **Software and Algorithms** | | |
| ImageJ | NIH | https://imagej.nih.gov/ij/ |
| ANY-maze 2.1 | Stoelting Co. | www.anymaze.co.uk |
| IMARIS 9.7.0 | Oxford Instruments | https://imaris.oxinst.com/ |
| Prism 6 | GraphPad | https://www.graphpad.com/scientific-software/prism/ |
| Zen Blue 2.3 | Zeiss | N/A |
| Zen Lite | Zeiss | https://www.zeiss.com/microscopy/int/products/microscope-software/zen-lite.html |
| Huygens software | Scientific Volume Imaging | https://svi.nl/Huygens-Software |
| PATCHMASTER | HEKA Elektronik | https://www.heka.com/downloads/downloads_main.html#down_patchmaster |
| Clampex 10.4 | Molecular Devices | https://www.moleculardevices.com |
| Campfit 10.4 | Molecular Devices | https://www.moleculardevices.com |
| Coreldarw X3 | CorelDRAW | https://www.coreldraw.com |
| MiniAnalysis 6.0.9 | Synaptosoft | http://www.synaptosoft.com/ |
| **Others** | | |
| Digital Small Animal Stereotaxic frame | David Kopf instruments | Cat# Ultra Precise 962/963 |
| Micro Temperature Controller | Physitemp | MTC-1 |
| Picospritzer III | Parker Hannifin | N/A |
| Capillary Borosilicate Glass | Warner | G100F-4 |

| | | |
|---|---|---|
| 1.0mm OD, 0.58mm ID, 10 cm length, with filament (for stereotaxic injection) | Instruments | |
| Fear Conditioning System | Ugo Basile | N/A |
| Lecia Vibrating Blade Microtome | Leica | Cat# VT1000S |
| Digitizer | Molecular Devices | Cat# Digidata 1440A |
| MultiClamp 700B Microelectrode Amplifier | Molecular Devices | Cat# MULTICLAMP 700B |
| EPC9/2 amplifier | HEKA Electronik | N/A |
| Pipette Puller | Zeitz-Instrumente Vertriebs | N/A |
| Borosilicate Glass | Harvard Apparatus | Cat# GC 120F-10 |
| Axio Imager | Zeiss | N/A |
| gSTED laser scanning microscope | Leica | TCS SP8 STED |
| Confocal Microscope | Zeiss | LSM 780 |

**Materials and Methods**

**LEAD CONTACT AND MATERIALS AVAILABILITY**

Further information and requests for resources and reagents should be directed to and will be fulfilled by the Lead Contact, Cheng-Chang Lien. This study did not generate any unique reagents.

**EXPERIMENTAL MODELS AND SUBJECT DETAILS**

Wild-type C57BL/6J mice were obtained from Janvier labs. The Sst-IRES-Cre, Fos2A-iCreERT2 (TRAP2) mice and the Ai14 reporter mice were purchased from the Jackson Laboratory (stock #013044, #030323, #007908). Prkcd-Cre mice were obtained from Mutant Mouse Research and Resource Centers (MMRRC Stock no. 011559-UCD). Sst-IRES-Cre and Prkcd-Cre mice were crossed to Ai14 mice for use in many experiments described in this study. All mice were bred onto the C57BL/6J genetic background. Mice with both sexes (postnatal 7-13 weeks) were used for the experiments. The mice had *ad libitum* access to food and water, and were group-housed on a 12-hour light/dark cycle. Mice were singly housed, transported, and handled once daily for 4 days prior to experiments. All experiments were performed during the light phase. Animal experiments were performed according to standard ethical guidelines and approved by the Danish National Animal Experiment Committee (Permission No. 2017−15−0201−01201), Animal Care and Use Committee of the National Yang Ming Chiao Tung University (Permission No. 1080317), and the Austrian Animal Experimentation Ethics Board (Permission No. 2020-0.547.574).

### METHOD DETAILS

#### Viral vectors

To conduct chemogenetic experiments on TRAP2 mice, we injected a viral vector ssAAV-2/1-hDlx-HBB-chI-dlox-HA_hKORD_mCyRFP1(rev)-dlox-WPRE-hGHp(A) ($6\times10^{12}$ vector genomes/mL, v326-1, VVF Zurich, Switzerland) to express inhibitory designer receptors exclusively activated by designer drugs (iDREADDs) on TRAPed neurons. For anterograde tracing experiments, we used a viral vector ssAAV2/1-EF1a-DIO-hChR2(HI34R)-mCherry ($5.1\text{x}10^{12}$ GC/ml, Addgene #20297).

#### Stereotaxic injection

Mice (Sst-IRES-Cre, n = 10; TRAP2, n = 31, 6-10 weeks old) were anesthetized using a mix of 0.05 mg/ml of Fentanyl (0.05 mg/kg, 154 007007, Hameln), 5 mg/ml of Midazolam (5 mg/kg, 002124, Hameln, 002124) and 1 mg/ml of Medetomidine (0.5 mg/kg, VM Pharma, 087896) (FMM). Mice were placed on a stereotaxic frame (Kopf instruments, CA, USA) and a homeothermic pad (MTC-1, Physitemp) was placed below the mice to maintain their body temperature at a constant 34-36 °C. After securing the head with ear bars, 70% ethanol was used to sterilize the surgical area along with Xylocaine (2%) as a local anesthesia and the eyes were protected with viscotears gel (Alcon). Analgesia (Buprenorphine, 0.1 mg/kg, Indivior) was administrated 30 minutes before end of surgery. A midline scalp incision (~0.8 cm) was made with scissors and the skin was pulled aside to expose the skull. Small craniotomies were made to target CeL bilaterally (coordinates from bregma: AP: −1.34 mm, ML: ±2.88 mm, DV: −4.5 mm) by drilling small holes using a high-speed drill (Foredom Electric Co, K.1070-22). The coordinates were normalized to a bregma-lambda distance of 4.21 mm. The viruses were injected through a glass capillary (Harvard Apparatus) by pulses of the Picospritzer III (Parker Hannifin), with 0.5 Hz pulse frequency and reach total 350 nl/hemisphere. The pipette was raised 0.1 mm above the injection sites for an additional 10 minutes to minimize the upward flow of viral solution and was slowly withdrawn. After viral injection, the incision was closed by suturing and mice were given an antidote mix of 0.4 mg/ml Naloxone (B. Braun, 115241; 1.2 mg/kg), 5 mg/ml Atipamelozone Hydrochloride (Revertor, Vibrac AG; 2.5 mg/kg) and 0.5 mg/ml Flumazenil (Sintetica; 0.5 mg/kg).

For anterograde tracing experiments, anaesthesia was induced with a combination of intraperitoneally injected Ketamine (80 mg/kg; Ketasol, AniMedica) and Xylazine (5 mg/kg; Xylasol, Animedica) and maintained with 2% Sevofluran (SEVOrane). The head was fixed on a stereotactic frame (Model 1900, Kopf Instruments) and ophthalmic ointment was applied to the eyes to avoid drying. Postoperative pain medication included administration of meloxicam (Metacam, Boehringer Ingelheim; 1 mg/ kg subcutaneously (S.C.)).

Sst-IRES-Cre mice (n = 2 females and n = 2 male) were unilaterally injected into the CeL in a volume of 0.2 µl using a glass pipette (tip diameter ~30 µm) connected to a Picospritzer III microinjection system (Parker Hannifin Corporation) at the following coordinates from bregma: for females: AP: -1.34 mm, ML: +3.0 mm, DV: -4.4 mm; for males: AP: -1.34 mm, ML: +3.0 mm, DV: -4.6 mm. Mice were allowed to recover for 3-4 weeks before perfusion to ensure adequate viral transduction. All animals were allowed to rest at least 4 weeks in the home cage before the behavioral experiments. We confirmed all viral injection sites through *post hoc* histology examination.

**Solutions and drugs**

4-hydroxytamoxifen (4-OHT; Sigma, Cat# H6278) was dissolved in dimethyl sulfoxide (DMSO) by shaking for 15 minutes and stored at –20°C for up to several weeks. 4-OHT was prepared freshly before the experiment under chemical hood by dissolving in 2% TWEEN80 (Sigma, Cat# P1754) (diluted in saline solution (0.9%)) to reach a final concentration of 10 mg/ml. The 4-OHT injections were delivered intraperitoneally (I.P.) with an injectable concentration of 15 mg/kg. Salvinorin B (SALB, HelloBio, Cat# HB4887) was dissolved in 0.9% NaCl with 10% DMSO, and injected S.C. with the final concentration 10 mg/kg. Control experiments were conducted by vehicle (Veh) injection containing 10% DMSO in 0.9% NaCl solution. For the *ex vivo* slice recording, the following antagonists were added to the ACSF: 2 mM kynurenic acid (Sigma-Aldrich) to block AMPA and NMDA receptors for spontaneous IPSC recordings, and additional 1 µM tetrodotoxin (Tocris) applied to block voltage-gated $Na^+$ channels for miniature IPSC recordings.

**Behavioral tests**

Apparatus

Fear conditioning and retrieval tests were performed in chambers (17x17x25 cm) with transparent walls and a metal rod floor (context A; UGO BASILE, #46003). Grid floor rods were connected to a shock generator. The chambers were placed in a sound-attenuating box with ventilating fan, a dual (visible/I.R.) light, a speaker and an USB-camera (UGO BASILE, #46001).

Fear conditioning

All mice were handled by the experimenter 4 days before the experiment day (Day 0). Mice (8- to 12-week-old, C57BL/6J, n=32; Sst-IRES-Cre;Ai14, n=19; Sst-IRES-Cre, n=10; Prkcd-Cre;Ai14, n=7, TRAP2;Ai14, n = 5) were subjected to an auditory fear-conditioning paradigm consisting of: day 0, mice were placed into the conditioning chamber. After a 120 s acclimation period, an 80 dB, 4 kHz pure tone [conditioned stimulus (CS)] lasting 20 s was paired in the last 2 s with a 0.6 mA scrambled footshock [unconditioned stimulus (US)] five times (60 s inter-pairing interval). Control mice (Ctrl, Cue-only) were exposed to the CS-only. Mice were returned to the home cage after a 120 s no-stimulus consolidation period. Chambers were cleaned between each animal with either 70% ethanol or 1% acetic acid.

For TRAP2 mice (n = 58), 3 weeks after viral transfection, the mice were handled by the experimenter 4 days before Day 0. To reduce background c-Fos activity, mice were dark adapted 24 hours before fear conditioning. On Day 0, the TRAP2 mice were subjected to auditory fear conditioning as described above, then injected with 4-OHT (15 mg/kg, I.P.) immediately after the fear conditioning under dim light, and were returned to the home cage in the dark room for further 24 hours.

Fear retrieval were performed 24 hours or 7 days after the conditioning in a novel context with striped or checkered curved walls and flat floors. Mice were placed into the new context. After a 120-s acclimation period, there were 3 CS presentations (20 s, 60 s inter-CS interval). TRAP2 mice were injected (S.C.) with either SALB (10 mg/kg) or Veh control 30 minutes prior to the test session. 3 mice of the TRAP2-hKORD group in experiments shown in Figures 5A-C were excluded due to the incorrect injection sites.

TRAP2 mice with viral transfection (n = 4) were tested with two distinct auditory fear conditioning paradigms (Figures S3I-M). On day 0, TRAP2 mice were subjected to auditory fear conditioning (FC1, CS1: 4 kHz) and 4-OHT injection as described. On day 7, they were subjected to FC2 containing 5 pairings of a 20 s, 80 dB auditory cue (CS2: 12 kHz) co-terminating with a 2s, 0.6 mA footshock (US), and were perfused within 90 minutes after the FC2 acquisition.

TRAP2 mice (n = 23) were tested with the two associative fear learning paradigm with DREADD inhibition (Figures 5D-F, Figures S6A-C). The context for FC1 conditioning is a standardized commercial mouse cage (L*W*H: 18*18*25 cm, Ugo Basile, #46002). The context for FC1 retrieval is a plastic cylinder covered by checkboard pattern paper with a steel bottom (diameter 18 cm, height: 28 cm). The context for FC2 conditioning is modified from the original mouse cage by masking it with strip pattern or white wall forming a spindle shape and wiped with 0.1% acetic acid. The context for FC2 retrieval is a plastic cylinder covered by red paper, the steel bottom is covered with a thin layer of bedding material (0.5 cm), which is different from what is used in their home cage. The mice were handled for 4 days prior to both fear conditioning and retrieval test. All behavioural experiments were conducted in the end of light cycle (ZT10-12). On day 0, TRAP2 mice were subjected to auditory fear conditioning FC1 (5 x CS1 + US, CS1: 4 kHz) and 4-OHT injection as described. On day 7, they were tested for FC1 memory retrieval. For experiments shown in Figures S6A-C, on day 7, the mice were injected with SALB (10 mg/kg, S.C.) 30 minutes prior to the FC1 test session. On day 8, they were subjected to FC2 (5 x CS2 + US, CS2, 12 kHz). On day 15, TRAP2 mice were injected (S.C.) with either SALB (10 mg/kg) or Veh control 30 minutes prior to FC2 retrieval and were perfused within 90 minutes after retrieval. Two mice of the TRAP2-hKORD group in the experiment shown in Figures 5D-F and one mouse in the experiment shown in Figures S6A-C were excluded due to the incorrect injection sites. The freezing behavior, defined as the absence of movement except respiration for at least 1 s, was manually scored from video recordings (ANYmaze 2.1, 30 frames/s).

Open field test (OFT)

The OFT was performed in a plastic and square bright-field chamber (45 × 45 × 45 cm). The center zone was defined as 22.5 cm × 22.5 cm in the center of the arena, the remaining part of the arena was defined as the margin zone. TRAP2 mice were injected (S.C.) with either SALB (10 mg/kg) or Veh control 30 minutes prior to the test. Mice were placed in the center of the arena at the beginning of the experiment and then allowed to explore freely for 10 minutes. The total travel distance and time in the center zone were analyzed based on the video recordings (ANYmaze 2.1, 30 frames/s). The chamber was cleaned with 70% ethanol between trials.

**Slice preparation and patch-clamp recording**

Mice were anesthetized with isoflurane and decapitated by appropriately trained researchers within 90 minutes after the fear memory test. Brains were removed and 300 µm-thick coronal sections were prepared by a vibratome (Leica) using ice-cold sucrose saline containing (in mM): 87 NaCl, 25 $NaHCO_3$, 1.25 $NaH_2PO_4$, 2.5 KCl, 10 glucose, 75 sucrose, 0.5 $CaCl_2$, and 7 MgCl2. Slices were recovered in oxygenated (95% $O_2$ and 5% $CO_2$) sucrose saline containing chamber at 34 ˚C for 30 minutes and kept at room temperature (RT) until used. During the experiment, slices were transferred to a submerged chamber and perfused with oxygenated artificial cerebrospinal fluid (ACSF) containing (in mM): 125 NaCl, 25 $NaHCO_3$, 1.25 $NaH_2PO_4$, 2.5 KCl, 25 glucose, 2 $CaCl_2$, and 1 $MgCl_2$, at RT. The red fluorescence and neurons in the CeL were visually confirmed and selected for recordings under infrared differential interference contrast (IR-DIC) CCD camera (Hamamatsu).

Whole-cell patch-clamp recordings were made with a Multiclamp 700B amplifier (Molecular Devices, Sunnyvale, CA, USA) or a HEKA amplifier (EPC9/2 amplifier HEKA Electronik, Lambrecht, Germany; Pulse software). Recording electrodes (2–5 MΩ) were pulled from borosilicate glasses (outer diameter, 1.5 mm; 0.32 mm wall thickness; Harvard Apparatus) and filled with chloride-rich internal solution containing (in mM): 144 KCl, 0.2 EGTA, 4 MgATP, 10 HEPES, 7 $Na_2$-phosphocreatine, 0.1 GTP, and 0.2 % biocytin (wt/vol, Thermo Fisher Scientific, Waltham, MA, USA) with pH adjusted to 7.3 with KOH. The series resistance (Rs) was compensated to 70-80% in the voltage-clamp configuration. The data were discarded if Rs > 20 MΩ or Rs change > 20 % throughout the entire recording. Signals were low-pass filtered at 4 kHz (four-pole Bessel) and sampled at 10 kHz using a digitizer (Digidata 1440A; Molecular Devices).

**Immunohistochemistry**

To identify the biocytin-filled neurons, brain slices were fixed overnight with 4% paraformaldehyde (PFA) (wt/vol) in phosphate-buffered saline (PBS). After washing with PBS 3 times, 0.3% Triton X-100 (vol/vol; USB Co., Cleveland, OH, USA) was added for 30 min. Then slices were incubated with streptavidin-conjugated Alexa Fluor 488/594 (1:400; Thermo Fisher Scientific, Waltham, MA, USA) in PBS containing 0.3% Triton X-100 (vol/vol; USB Co., Cleveland, OH, USA) and 2% normal goat serum (NGS, Vector Laboratories, Burlin-game, CA, USA) at 4°C overnight. After washing 6 times with PBS, slices were mounted onto slides using Vectashield mounting medium containing 4,6-diamidino-2-phenylindole (DAPI, H-1200, Vector Laboratories, Burlin-game, CA, USA).

For the quantification of TRAPed cells and c-Fos expression, within 90 minutes after the cued fear memory test session, TRAP2-hKORD mice were anesthetized by FMM injection (I.P.) and were perfused with ice-cold PBS plus heparin (50mg/ml) followed by 4% PFA. Fixed brains were removed and underwent post-fixation in 4% PFA for maximum 30 minutes and were stored in 0.1M PB. Brains were sliced into 50 um coronal sections using a vibratome (Leica 1000 S). Collected brain slices were washed with TBS and underwent a two-step blockade: first against Mouse-on-Mouse blocking reagent (M.o.M, 1:27.7 in TBS) for 1 hour at RT, and then against normal horse serum (NHS) (containing 10% NHS and 0.1% Triton-X100 in TBS) for 1 hour at RT. Then slices were incubated overnight with primary antibody: anti-c-Fos (1:2000), anti-Somatostatin (1:250), anti-PKC-δ (1:600), anti-red fluorescence protein (anti-RFP, 1:500) with 2% NHS and 0.1% Triton-X100 in TBS for 48-72 hours at 4°C. After washing trice with TBS, slices were incubated with secondary antibodies: Alexa 488 (1:1000), Alexa 568 (1:1000), Cy3 (1:500), Cy5 (1:500), with 2% NHS and 0.1% Triton-X100 in TBS for 24 hours at 4°C. After washing with TBS, slices were mounted onto slides with

DAPI containing mounting medium (Vectashield/FluoromountG). Images were taken as z-stacks by a confocal microscope (LSM 780) by 2 µm step size. Measures of c-Fos+ cell proportion by area were manually counted from these images with IMARIS 9.7.0 software (Oxford Instruments, Bitplane, Zurich, Switzerland) by experimenters unaware of the treatment condition.

Immunofluorescence for the anterograde tracing experiments (Sst-IRES-Cre mice: n = 2 females and n = 2 males; TRAP2 mice: n= 6 females) were carried out according to previously published procedures with minor modifications (Sreepathi and Ferraguti, 2012). Mice were then deeply anaesthetized with thiopental sodium (150 mg/kg, I.P.) and transcardially perfused with a fixative made of 4% paraformaldehyde, 15% picric acid in 0.1 M phosphate-buffer (PB), pH 7.2-7.4. Brains were extracted from the skull and cut into 50 µm thick coronal sections using a vibratome (Leica Microsystems VT1000S, Vienna, Austria).

The fluorescence signal for mCherry was enhanced using a goat primary antibody against RFP (Rockland, cat. no. 200-101-379S; diluted 1:1000), which also reacts with mCherry, and a donkey anti-rabbit secondary antibody conjugated with Cy3 (Jackson ImmunoRes., cat. no. 705-165-147; diluted 1:500). Primary and secondary antibodies were prepared in 2% normal horse serum (NHS) and 0.1%Triton X-100 in TBS (TBS-T). Sections were then mounted onto gelatin-coated slides and coverslipped with ProLong Diamond antifade mountant (ThermoFischer Scientific).

Low magnification images were acquired using an epifluorescence microscope (Axio Imager, Carl Zeiss, Oberkochen, Germany) and the Openlab software (Version 5.5.0). For higher magnification aLeica TCS SP8 gSTED laser scanning microscope (Leica Microsystem GmbH, Germany) with a 63x/1.3 objective was used. Raw images were deconvolved using the Huygens software (Scientific Volume Imaging, Hilversum, The Netherlands). Image processing was performed using the IMARIS 9.7.0 software (Oxford Instruments, Bitplane, Zurich, Switzerland).

**Data analysis and statistics**

MiniAnalysis (6.0.9 Synaptosoft, Fort Lee, NJ, USA) was used to detect and analyze sIPSCs and mIPSC events. The SALB induced membrane potential and input resistance changes were analyzed using Clampfit version 10.4 (Molecular Devices). For fear conditioning, the freezing time is defined as a lack of movement associated with a crouching (>0.9 s) and the video is scored by Any-maze software (Stoelting) and manually checked by an experimenter blinded to the test groups. We used IMARIS software (Oxford Instruments) for the quantification of cell number and the molecular identity. Statistical tests were performed and plotted using Prism 6.0 (GraphPad Software, La Jolla, CA, USA). Statistical significance was tested by the Mann-Whitney test, Wilcoxon signed rank test, non-zero null test for linear regression, Kruskal-Wallis test (with Dunn's *post hoc* test), or two-way ANOVA at the significance level ($p$) indicated. Data were presented as mean ± standard error of mean (SEM). Significance levels were set at $^*p < 0.05$.
